# Structure of the cell-binding component of the *Clostridium difficile* binary toxin reveals a novel macromolecular assembly

**DOI:** 10.1101/833699

**Authors:** Xingjian Xu, Raquel Godoy-Ruiz, Kaylin A. Adipietro, Christopher Peralta, Danya Ben-Hail, Kristen M. Varney, Mary E. Cook, Braden M. Roth, Paul T. Wilder, Thomas Cleveland, Alexander Grishaev, Heather M. Neu, Sarah Michel, Wenbo Yu, Dorothy Beckett, Richard R. Rustandi, Catherine Lancaster, John W. Loughney, Adam Kristopeit, Sianny Christanti, Jessica W. Olson, Alex D. MacKerell, Amedee des Georges, Edwin Pozharski, David J. Weber

## Abstract

Targeting *Clostridium difficile* infection (CDI) is challenging because treatment options are limited, and high recurrence rates are common. One reason for this is that hypervirulent CDI often has a binary toxin termed the *C. difficile* toxin (CDT), in addition to the enterotoxins TsdA and TsdB. CDT has an enzymatic component, termed CDTa, and a pore-forming or delivery subunit termed CDTb. CDTb was characterized here using a combination of single particle cryoEM, X-ray crystallography, NMR, and other biophysical methods. In the absence of CDTa, two novel di-heptamer structures for <u>a</u>ctivated CDTb (aCDTb; 1.0 MDa) were solved at atomic resolution including a symmetric (^Sym^CDTb; 3.14 Å) and an asymmetric form (^Asym^CDTb; 2.84 Å). Roles played by two receptor-binding domains of aCDTb were of particular interest since RBD1 lacks sequence homology to any other known toxin, and the RBD2 domain is completely absent in other well-studied heptameric toxins (i.e. anthrax). For ^Asym^CDTb, a novel Ca^2+^ binding site was discovered in RBD1 that is important for its stability, and RBD2 was found to be critical for host cell toxicity and the novel di-heptamer fold for both forms of aCDTb. Together, these studies represent a starting point for structure-based drug-discovery strategies to targeting CDT in the most severe strains of CDI.

**SIGNIFICANCE STATEMENT:** There is a high burden from *C. difficile* infection (CDI) throughout the world, and the Center for Disease Control (CDC) reports more than 500,000 cases annually in the United States, resulting in an estimated 15,000 deaths. In addition to the large clostridial toxins, TcdA/TcdB, a third *C. difficile* binary toxin (CDT) is associated with the most serious outbreaks of drug resistant CDI in the 21^st^ century. Here, structural biology and biophysical approaches were used to characterize the cell binding component of CDT, termed CDTb, at atomic resolution. Surprisingly, two novel structures were solved from a single sample that help to explain the molecular underpinnings of *C. difficile* toxicity. These structures will also be important for targeting this human pathogen via structure-based therapeutic design methods.

## INTRODUCTION

Symbiotic microbiota in the gut typically prevent *C. difficile* colonization in healthy individuals, but as protective bacteria are reduced by common antibiotic treatment(s), cancer therapy and/or by other means, then CDI becomes a much higher health risk (1, 2). Upon diagnosis, it is critical to cease delivery of problematic antibiotics, particularly those prone to select for hypervirulent CDI strains (i.e. fluoroquinolones, clindamycin, cephalosporins) (3, 4), and then clear the infection with a limited choice of antibiotics that can sometimes provide efficacy including metronidazole, vancomycin and/or fidaxomicin (1, 5). However, continued resistance to antibiotics and overwhelming levels of toxin production by the *C. difficile* bacteria can severely limit such a clinical approach. Other options for patients having severe CDI are colonoscopy or experimental procedures such as a fecal microbiota transplant (FMT), but these treatment options can have severe drawbacks (1, 6). Consequently, novel therapies are needed, particularly for recurrent CDI and for cases associated with hypervirulent strains of *C. difficile* (i.e. BI, NAP1, 027, 078, and others) (1, 5, 7–9).

Antibiotic and anti-toxin combination therapy is often an effective clinical approach for toxin producing infections (10), so this strategy is under development for treating *C. difficile* infection (CDI). While therapeutic options are becoming available to target the large clostridial toxins (LCTs), TcdA/TcdB (11), there is nothing approved by the FDA to target the *C. difficile* toxin (CDT) or the “binary toxin” (12). Other evidence demonstrating an urgency to develop CDT anti-toxins include (i) patients with CDT-containing strains of CDI show heightened disease severity and reoccurrence (13–16); (ii) strains of *C. difficile* having only CDT and not TcdA/TcdB (A^−^B^−^CDT^+^) retain virulence and present as CDI in the clinic (16, 17), and (iii) an immunological response in hamsters to a vaccine targeting TcdA/TcdB and CDT showed much higher efficacy towards challenges from a hypervirulent strain of CDI (i.e. NAP1) than a vaccine derived only from TcdA/TcdB antigens (12, 18). Therefore, to address this unmet medical need, studies of the structure, function and inhibition of CDT are paramount to identifying its vulnerabilities and for developing novel treatments to improve patient outcomes for the most severe cases of CDI.

CDT is a binary toxin that has an enzymatic subunit, CDTa (47.4 kDa), with ribosyltransferase activity, and a pore-forming delivery subunit, termed CDTb (99 kDa) (15, 19–23). Prior to cellular entry via endosomes (24–27), CDT associates with host cell receptor(s), such as the lipolysis-stimulated lipoprotein receptor (LSR) and/or CD44 (28–31). Based on studies with other binary toxins, it was suggested that the low pH in endosomes triggers CDTa translocation into the cytoplasm, via the cell-binding and pore-forming entity, CDTb, but a detailed molecular mechanism for this process remains unknown (32–41). Once the CDTa enzyme is delivered into the host cell cytoplasm, ADP-ribosylation of G-actin occurs catalytically at Arg-177 (42). ADP-ribosylated G-actin then leads to F-actin filament dissociation (43), destruction of the cytoskeleton, increased microtubule protrusions, accelerated bacterial adhesion, and a *“death spiral*” for host cells (44–46). In this study, a combination of biophysical and structural biology methods was used to define the molecular structure of activated CDTb (i.e. termed aCDTb). The roles played by the two receptor-binding domains of aCDTb (RBD1, RBD2) were of particular interest in this study. RBD1 lacks sequence homology to any other known toxin and was found to have a novel Ca^2+^-binding site. The other receptor binding domain, RBD2, is at the C-terminus of aCDTb, and it is not present in other members of this toxin family. Importantly, RBD2 was shown to be critical for establishing the novel di-heptamer macromolecular assembly in aCDTb that is necessary for host cell toxicity. Together, these and other regions of aCDTb can now be considered in future structure-based drug design strategies.

## RESULTS AND DISCUSSION

### Structural and biophysical characterization of aCDTb

For studies of aCDTb, pro-CDTb (residues 1-876) was overexpressed in baculovirus-infected insect cells and purified to homogeneity. Active CDTb (aCDTb; residues 212-876) was generated via limited proteolysis with chymotrypsin to remove the signaling peptide (SP) and the activation domain (AD, residues 43-211) with hydrolysis confirmed to be between M211 and S212 by mass spectroscopy, as previously described (12). The aCDTb protein was purified to homogeneity (>99%) and shown to be fully active in *Vero* cell killing assays using catalytic amounts of aCDTa (^CDTa^TC_50_=110±10 pM; Fig. 1) and an optimal CDTa to CDTb ratio of 1:7 was observed, as previously described (47).

**Figure 1.**
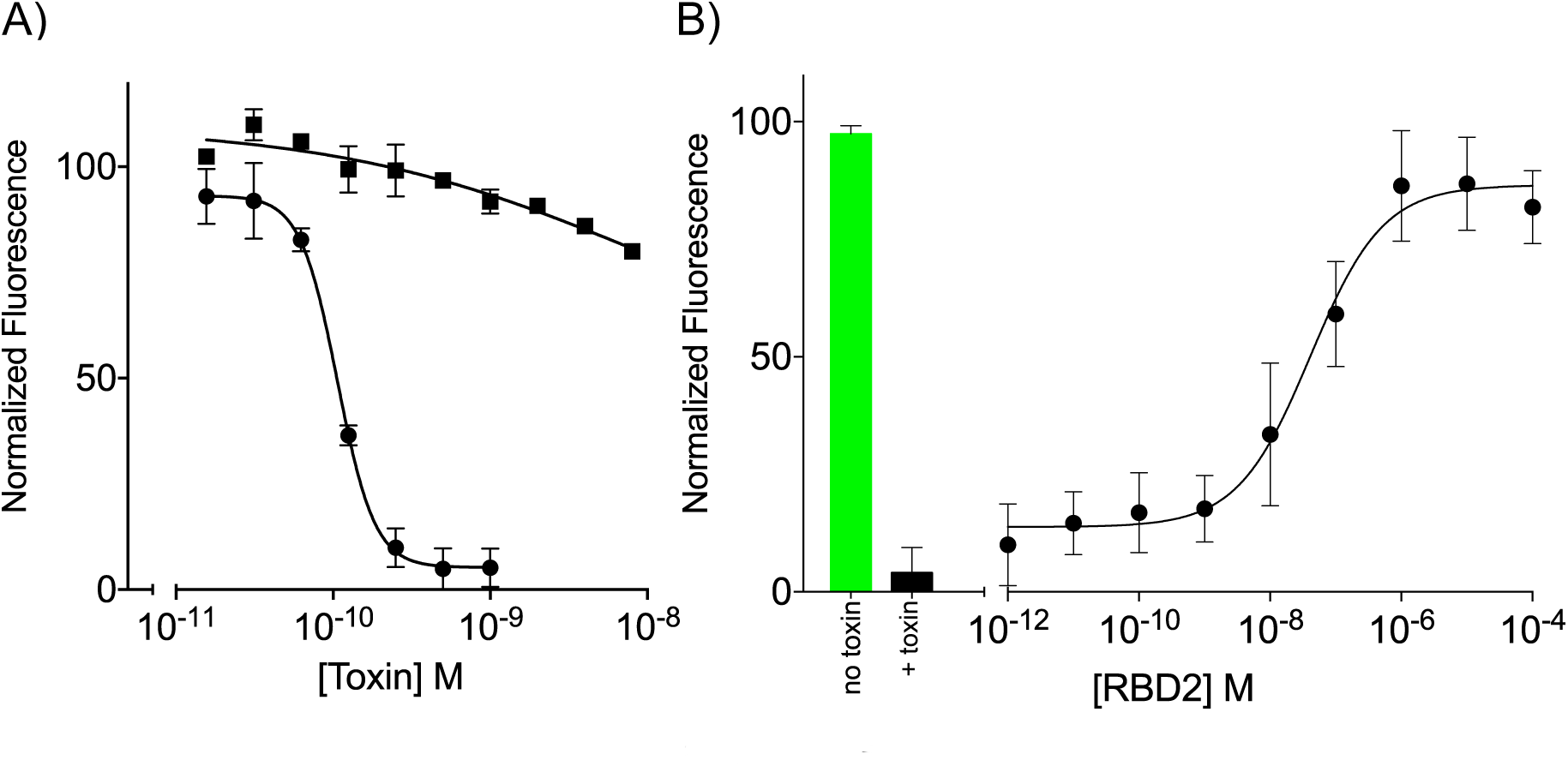
Functional studies of the receptor binding domains of CDTb (RBD1/RBD2). (**A**) Cellular toxicity upon the addition of CDTa to vero cells in the presence of aCDTb (□) or aCDTb^∆RBD2^ (n). The TC_50_ of CDTa is 150 ± 40 pM (n=8 independent experiments, ± S.D) when aCDTb is present. Little or no toxicity is observed when ^1m^CDTa is added vero cells with aCDTb^∆RBD2^ even at the highest concentrations (≥10 nM; n=3). For simplicity, the X-axis presented using the ^1m^CDTa concentration, but each experiment contains a 7x concentration of aCDTb or aCDTb^∆RBD2^, as previously described. (**B**) A representative experiment showing the effect of adding the RBD2 domain of CDTb (residues 757-876) into a vero cell toxicity assay with 500 pM binary toxin. These data illustrate that the isolated RBD2 domain inhibits cellular toxicity, as a dominant negative (IC_50_ = 20±10 nM; n=3) calculated using a four-parameter logistic regression analysis. All data are plotted versus the normalized fluorescence of Alexa Fluor 488 Phalloidin, which selectively labels F-actin filaments; an increase in toxicity causes depolymerization of actin, causing a decrease in fluorescent signal.

Sizing studies of aCDTb were completed using sedimentation velocity analytical ultracentrifugation (AUC) and size exclusion chromatography-multiangle light scattering (SEC-MALS) to determine its subunit stoichiometry. Surprisingly, rather being heptameric, as described for other cell-binding components of binary toxins (41), both methods showed monomeric aCDTb (75 kDa) to be the major species (95±2%) and a novel fourteen-subunit oligomer (1.0 MDa) detected at lower levels (<4-6%; Fig. 2A/2B). There was no evidence for heptameric aCDTb (< 0.1%). Consistent with SEC-MALS and AUC data, the presence of the fourteen-subunit aCDTb oligomer was validated using small angle X-ray scattering (SAXS) with its radius of gyration found to be 86±2 Å and a molecular weight of 1.0 ± 0.2 MDa (Fig. 2C). The interatomic distance probability distribution calculated from the SAXS scattering profile indicated that the aCDTb had a maximum particle dimension of 270 Å, and modeling these data with two dumbbell-like shapes of the di-heptamer markedly improved the quality of fit. Importantly, single-particle cryoelectron microscopy (cryoEM) studies were completed for aCDTb (**Figs. S1, S2**), in the absence of CDTa, and demonstrated unambiguously that the 14mer oligomerization state of aCDTb was the only higher molecular weight state observed, but interestingly, it had two unique structures including a symmetric (^Sym^CDTb) and an asymmetric (^Asym^CDTb) dimer of heptamers, which were solved at resolutions of 3.14 Å and 2.84 Å, respectively (Figs. 3–7). Likewise, crystals were obtained from the same aCDTb preparations and the availability of cryoEM models was essential to solving its structure by molecular replacement. The results of the X-ray studies further confirmed the dimer of heptamer stoichiometry for CDTb; however, only the ^Asym^CDTb was observed when the X-ray diffraction data were analyzed at 3.70 Å resolution (**Fig. S3**). These novel aCDTb structures will be important for continued delineation of the toxin’s mechanism of action as well as for future drug development efforts targeting the *C. difficile* binary toxin (CDT).

**Figure 2.**
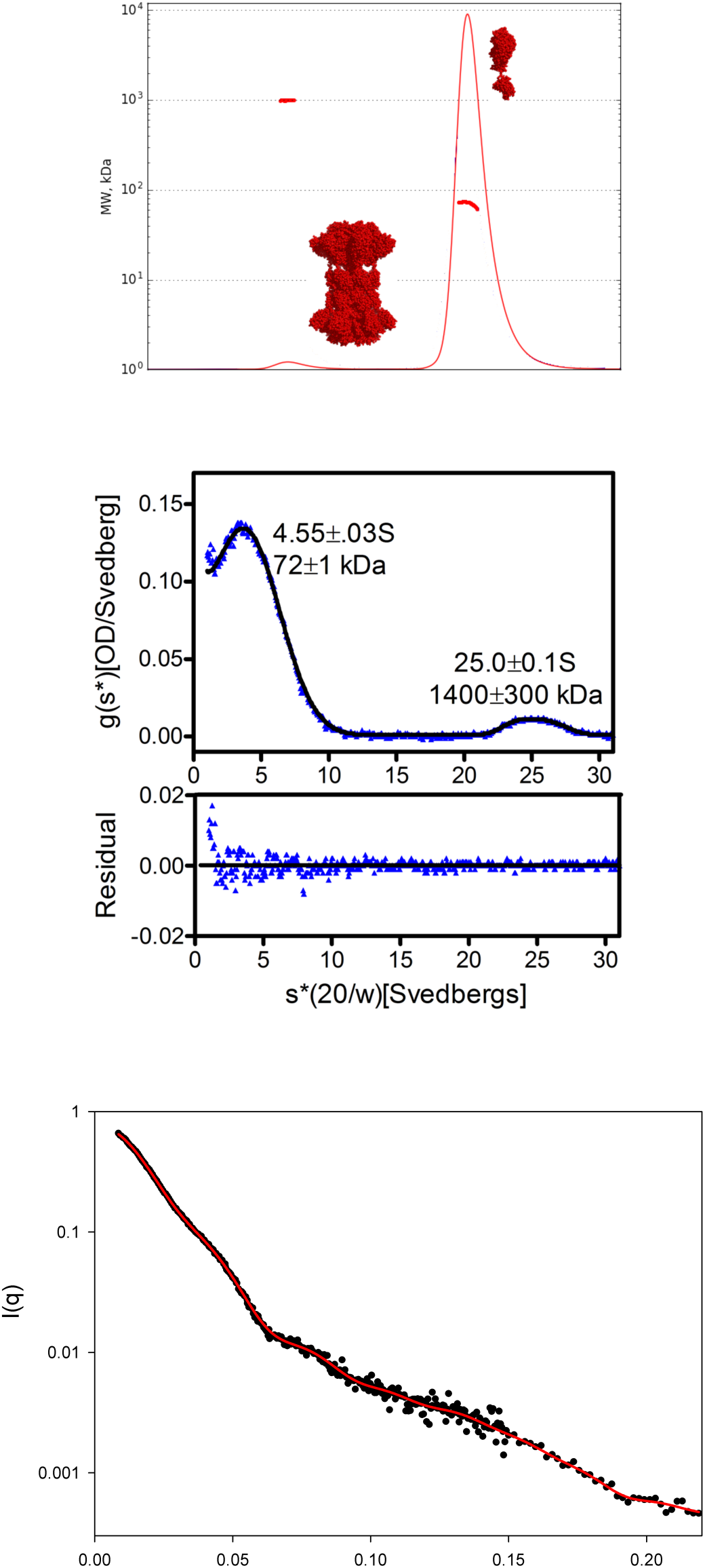
Biophysical studies of the oligomerization state of activated CDTb. **(A)** SEC-MALS trace for activated CDTb. Trace represents absorbance measurements; red dots are molecular weight estimates. Representative model structures are shown to designate the corresponding peaks. **(B)** Sedimentation velocity analysis of an 11 µM CDTb sample indicates that it is predominantly monomeric. Top Panel: The time-derivative distribution (blue triangles) and the best-fit of the data to a two species model (black line). Bottom Panel: Residuals of the fit to a two-species model. The errors in the sedimentation coefficients (s*(20/w) and the molecular weights represent the 95% confidence intervals. **(C)** Small angle X-ray scattering curve for activated CDTb. Experimental data (black dots) is shown and fitted with a model that included a mix of ^Asym^CDTb and ^Sym^CDTb at approximately 1:1 ratio.

**Figure 3.**
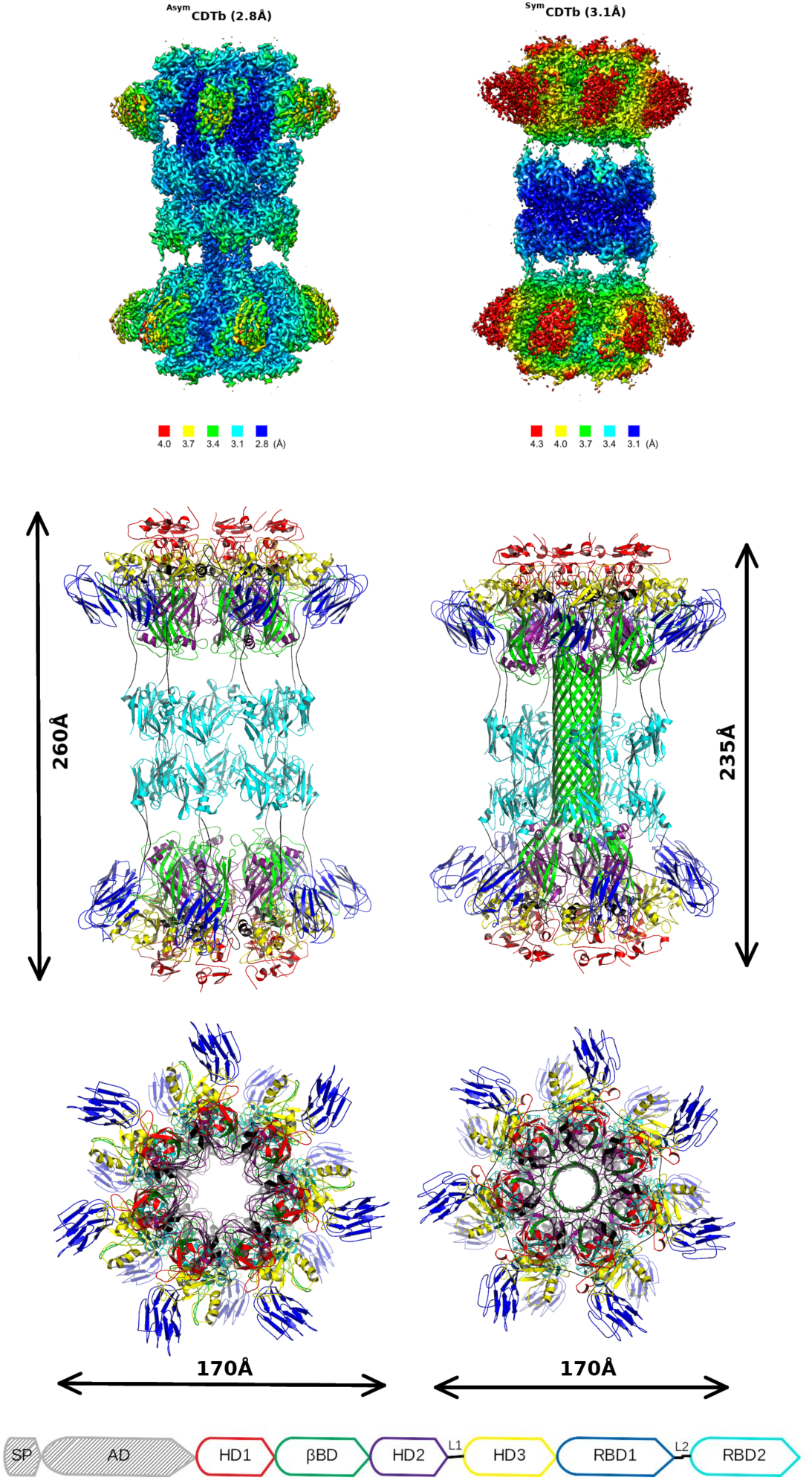
Structure of aCDTb. (**A**) Local resolution in structures of ^Asym^CDTb and ^Sym^CDTb conformations. Increased flexibility is observed in outer regions of the core heptamer, most pronounced for the RBD1 domain. (**B**) Overall structure of the aCDTb tetradecamer in ^Asym^CDTb and ^Sym^CDTb conformations. Color scheme is shown in domain diagram and both models are on the same scale, demonstrating slight shortening of the ^Asym^CDTb. Domains include a heptamerization domain (HD1; residues 212-297; Figs. S4, S5), a β-barrel domain (βBD; residues 298-401; Figs. S6, S6), a second heptamerization domain (HD2; residues 402-486; Figs. S8), a linker region (L1; residues 487-513; Figs. S6, S7), a third heptamerization domain (HD3; residues 514-615; Fig. S9), a receptor binding domain (RBD1; residues 616-744; Fig. S10), a second linker (L2; residues 745-756; S6, S7), and a second receptor binding domain (RBD2; residues 757-876; Fig. S11).

### ^Sym^CDTa and ^Asym^CDTb at atomic resolution

The X-ray and cryoEM structures of the cell binding and delivery component of the binary toxin, aCDTb, were examined in detail. Single particle cryoelectron microscopy (cryoEM) studies of aCDTb revealed two unique structures including a symmetric (^Sym^CDTb) and an asymmetric (^Asym^CDTb) form (Fig. 3), and the ^Asym^CDTb form was confirmed via X-ray crystallography (**Fig. S3**).

### The global folds of the two di-heptamer aCDTb structures

The heptamer units in the di-heptamers of CDTb assume two distinct forms. As shown (Figs. 3,4,5), an extended β-barrel resides in one of the heptamer units that resembles the low pH membrane inserted structure of the protective antigen (PA) cell-binding component of the anthrax toxin (48, 49) while the other lacks this structural motif and is more similar to the soluble form of the anthrax toxin. Nonetheless, while there are some similarities, the structures of both CDTa di-heptamers differs significantly from the heptameric assembly characteristic of the pore-forming component of the anthrax PA (Fig. 5) (41, 48, 50–52). Specifically, for ^Sym^CDTb, both heptamers of the di-heptamer are in a non-β-barrel form. The non-β-barrel/ non-β-barrel assembly of the two heptamers for ^Sym^CDTb is driven by a central donut-like structure formed by 14 copies of the 14 kDa C-terminal domain of CDTb (termed RBD2), which is absent in the anthrax PA (Fig. 5). Whereas, ^Asym^CDTb comprises a mixed non-β-barrel/β-barrel di-heptamer assembly, but again the two heptamer assemblies of this asymmetric form are still brought together as a di-heptamer by this unique RBD2-mediated mechanism. In ^Asym^CDTb, the 105 Å long β-barrel structure makes additional non-β-barrel/β-barrel interactions with the RBD2 domains (Fig. 7A) and shield several hydrophobic residues, which likely stabilizes ^Asym^CDTb prior to CDTa/receptor binding and/or insertion into the lipid membrane of host cells.

**Figure 4.**
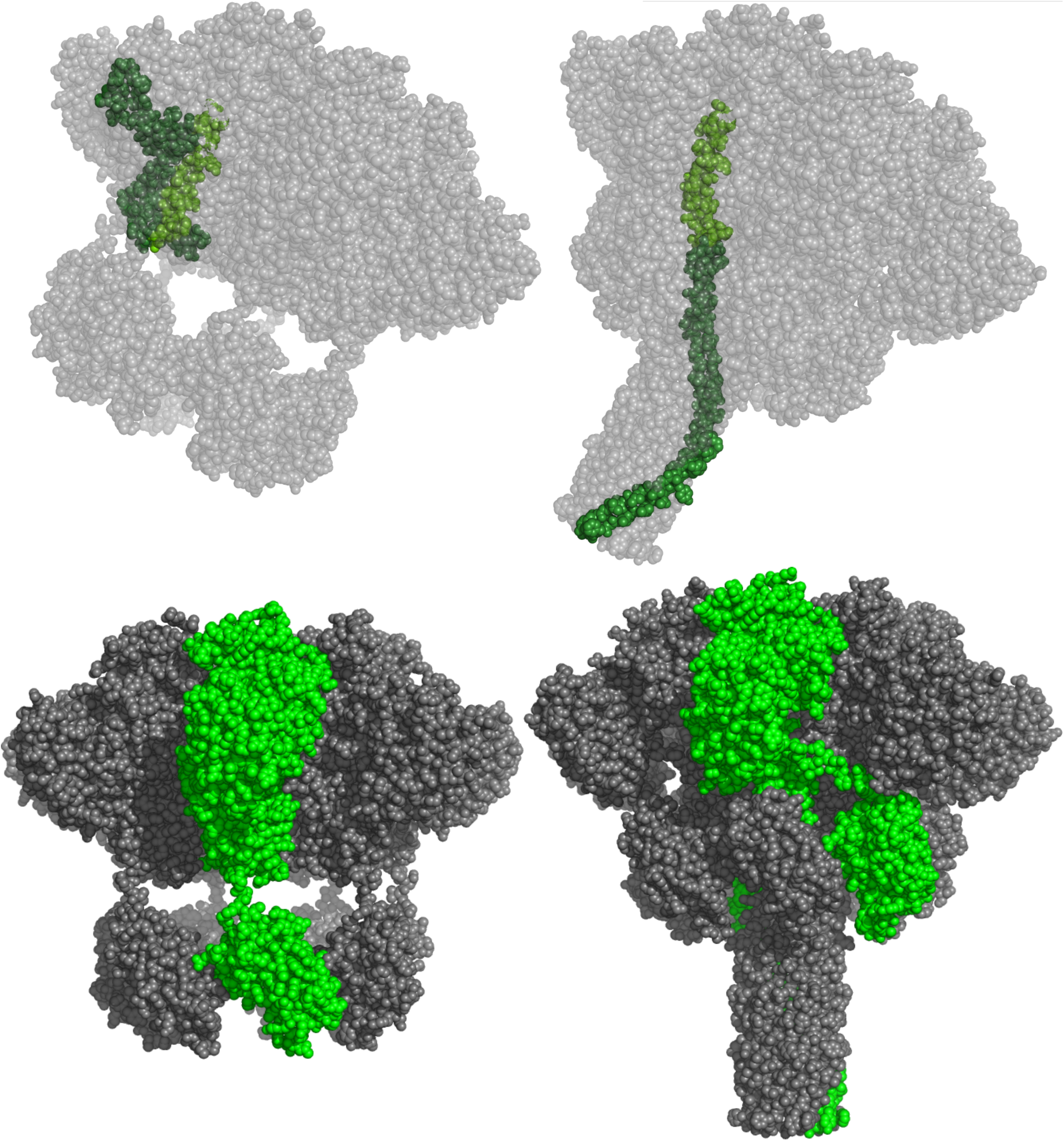
Large conformational differences when the two heptamer domains of ^Sym^CDTb without the beta barrel and ^Asym^CDTb having the beta barrel are compared. (**A**) Different packing of the β-barrel domain occurs in the two different heptamer conformations. For visualization, the single chain of the β-barrel domain is highlighted in lighter green. (**B**) The RBD2 domain donut assembly is located differently in the two different heptamer conformations shifts. For clarity, a single polypeptide chain is highlighted in green to show the varied arrangement of the RBD2 domains.

**Figure 5.**
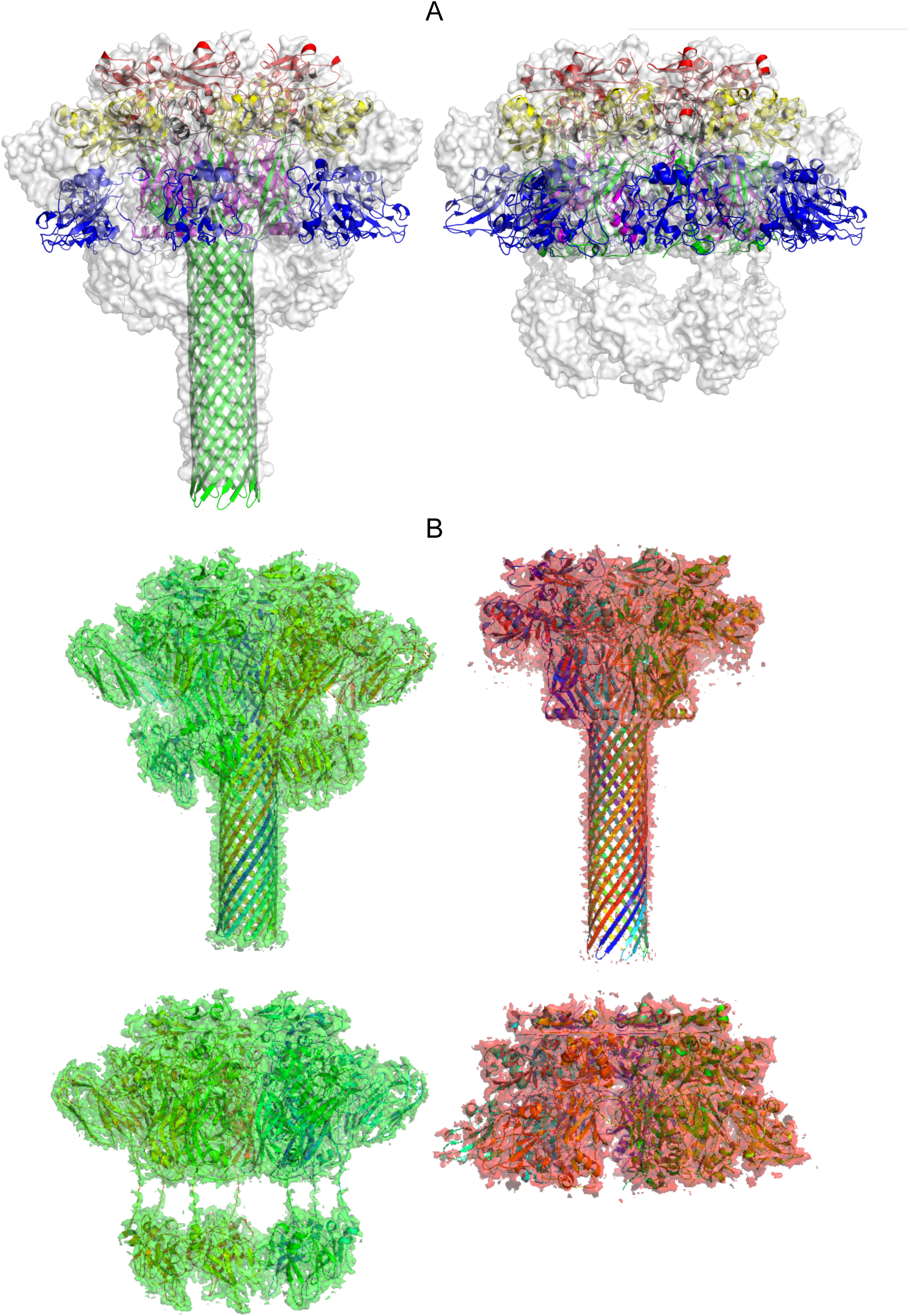
Comparison of the “β-barrel” containing heptamer of ^Asym^CDTb to the analogous heptamer from the protective antigen (PA) of the anthrax toxin. (**A**). Heptamers from PA of anthrax toxin are superimposed with electron density from the “β-barrel heptamer” observed in the ^Asym^CDTb di-heptamer structure. The receptor binding domain in “β-barrel form” of the PA from anthrax toxin were not modeled in the corresponding cryoEM model and are placed here using alignment with the soluble form of the toxin. (**B**) Structural comparison of heptameric forms A (top) and B (bottom) from ^Asym^CDTb (green) and anthrax toxin (red). CryoEM densities are shown for all molecules except anthrax toxin form B for which the 2Fo-Fc map is shown and derived from the corresponding crystal structure.

**Figure 6.**
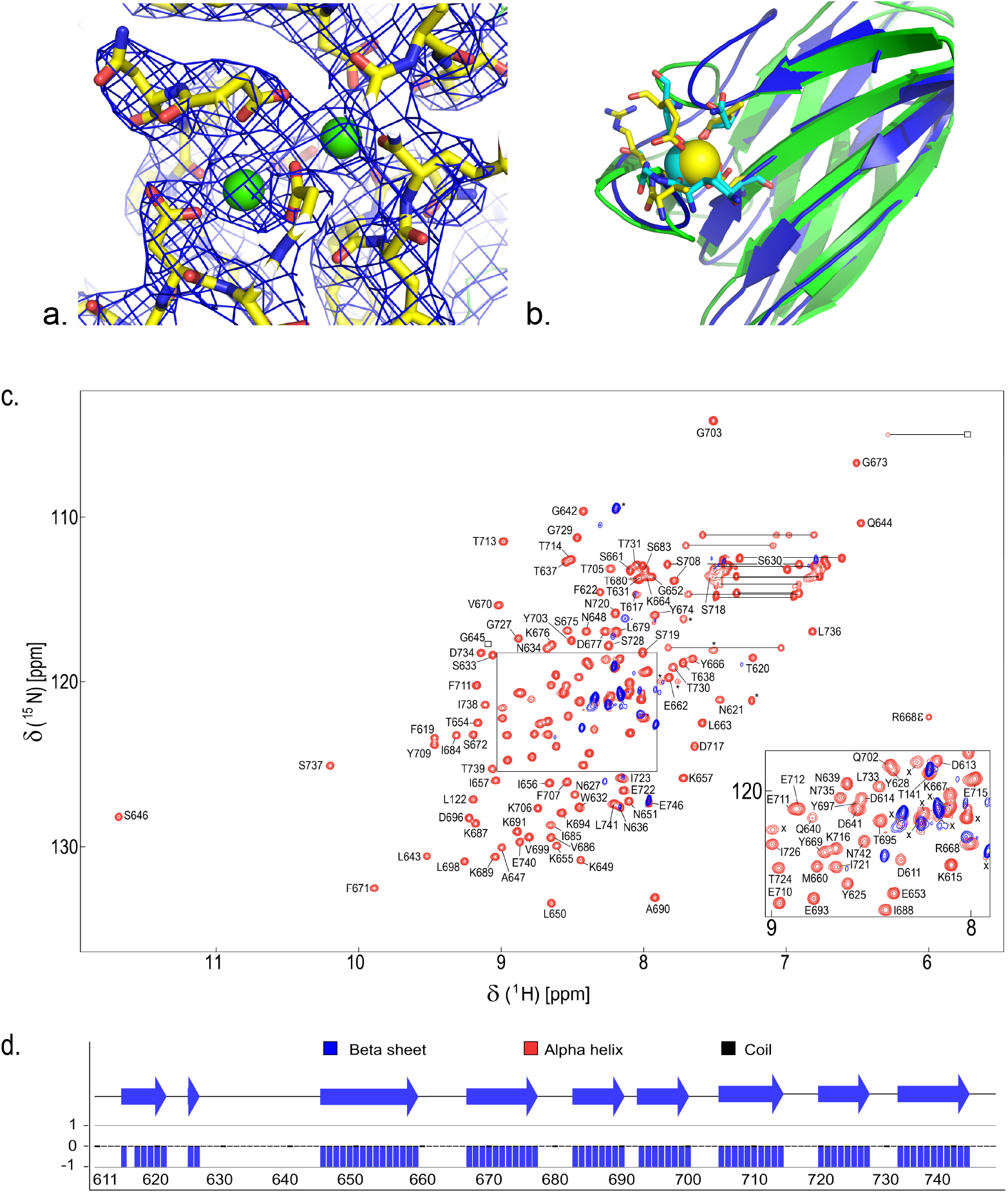
Detailed structural features of the aCDTb receptor binding domains. (**A**) Dual calcium binding site located in the N-terminal region of the protein. The coulomb potential maps (i.e. cryoEM density maps) in both ^Sym^CDTb and ^Asym^CDTb resolved two Ca^2+^ ions bound (Ca1, Ca2) with Ca1 oxygen ligands from D222/D224/E231/D273/N260(C=O)/E263(C=O), and Ca^2+^ ligands from D220/D222/D224/E321/D228/I226(C=O). (**B**) Calcium-binding site located in the β-sandwich domain of RBD1. The RBD1 domain of aCDTb is shown in blue, superimposed with the structure of the β-sandwich from *C. thermocellum* xylanase Xyn10B used here as an example of Ca^2+^-binding CBM domain. (**C**) Calcium is required for stability of the isolated RBD1 domain. The ^1^H,^15^N-HSQC spectrum of the RBD1 domain is illustrated in the absence (blue) and presence (red) of 6 mM CaCl_2_. A large number of the correlations, in the absence of Ca^2+^ (blue) were absent due to exchange-broadening or very strong (marked by X) consistent with this construct being “unfolded” in the absence of Ca^2+^. Upon Ca^2+^addition, the backbone and sidechain (i.e. for R668 epsilon) correlations appeared and were highly dispersed, consistent with the RBD1 domain folding in a Ca^2+^-dependent manner. Labeled are resonance assignments for ^1^H-^15^N correlations (in red) that are fully correlated with their corresponding ^13^C_alpha_ and ^13^C_beta_ chemical shift values, along with 96% of C’ shifts, and 93% of the sidechain shift values from triple resonance heteronuclear NMR data (SI). Seven other correlations were not assigned (marked with *) due to a complete lack of interresidue correlations in the triple-resonance NMR spectra; perhaps some of these unassigned correlations arise from the 6-residue His-Tag used for purifying this domain. Nine other observable correlations (red; labeled with an X) were not assigned, even in the presence of Ca^2+^, and remain disordered based on their narrow line shape and high intensity. Similarly, twenty-nine ^15^N-^1^H correlations (in blue) for residues of RBD1, in the absence of Ca^2+^, were not be readily assigned due to their intrinsically disordered state. (D) The predicted secondary structure of RBD1 in the presence of Ca^2+^ is predominantly beta strands and consistent with that of the RBD1 cryoEM structures (Fig. 3; Fig. S10), and is comprised of nine beta strands spanning residues: K615-N621; Y625-N626; G645-P659; K667-D677; S683-A690; E693-P700; T705-T714; N720-G727, and Y732-N742.

**Figure 7.**
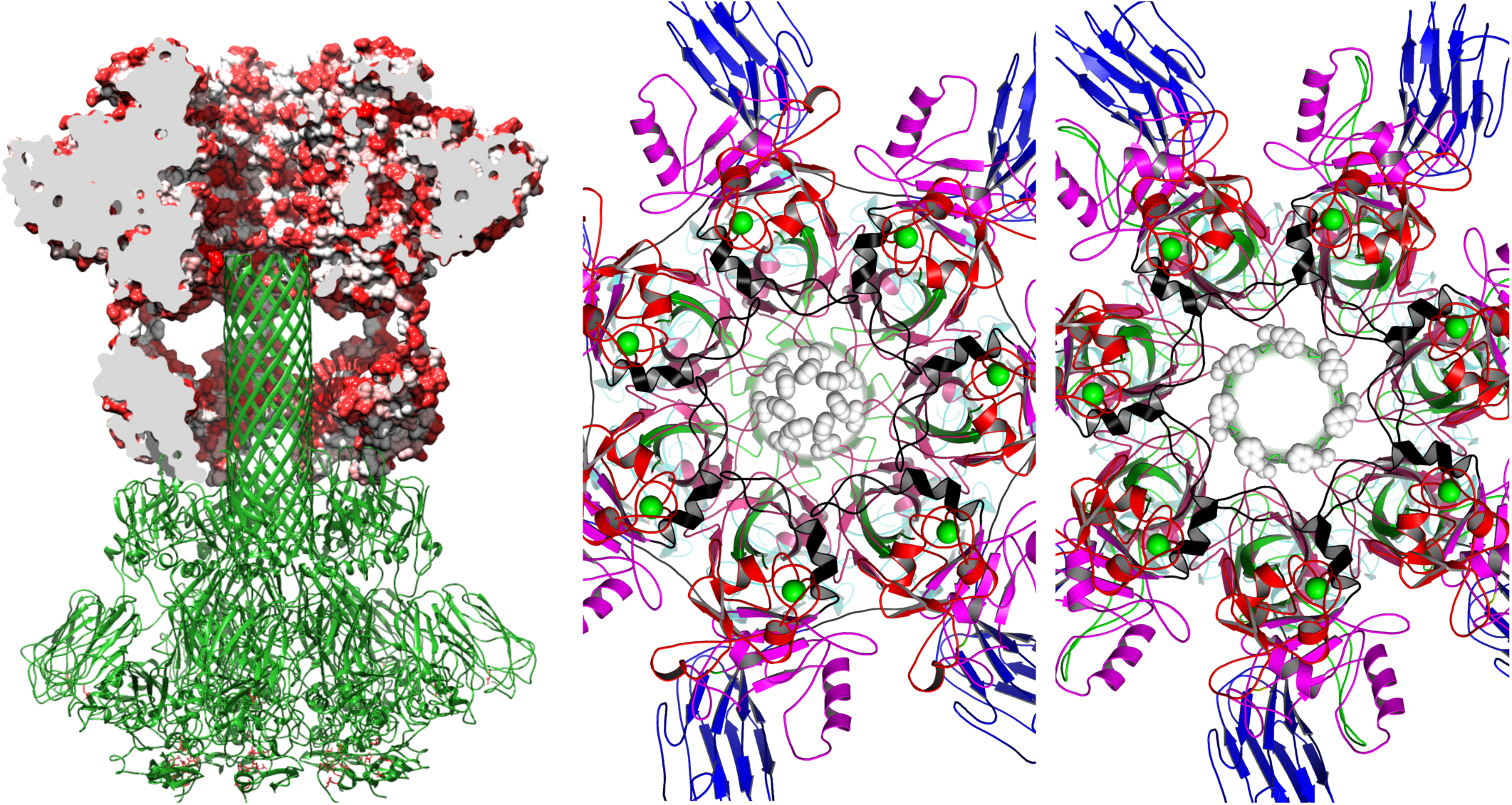
Large scale structural features of the aCDTb. (**A**) Cross heptamer hydrophobic interactions between the tip of the βBD in barrel conformation and HD2 for ^Asym^CDTb. (**B/C**) Changes in the size of the pore formed by φ-gate residues (F455) in two heptameric forms of the aCDTb – ^Asym^CDTb (panel B) and ^Sym^CDTb (panel C).

### Specific domain structures within ^Asym^CDTb and ^Sym^CDTb

For more detailed comparisons, delineation of domains of aCDTb are based on homologous domains from heptameric toxins (Fig. 3). These include a heptamerization domain (HD1; residues 212-297; Figs. S4, S5), the β-barrel domain (βBD; residues 298-401; Figs. S6, S7), a second heptamerization domain (HD2; residues 402-486; Fig. S8), a linker region (L1; residues 487-513; Figs. S6, S7), a third heptamerization domain (HD3; residues 514-615; Fig. S9), a receptor binding domain (RBD1; residues 616-744; Fig. S10), a second linker (L2; residues 745-756; Figs. S6, S7), and a second receptor binding domain (RBD2; residues 757-876; Fig. S11). It is important to point out that RBD1 is not homologous to any other binary toxin, and when aligned, no other toxin was found to have an RBD2 domain. On the other hand, as for other heptameric pore-forming toxins, HD1, HD2, HD3, and RBD1 comprise a large number of the interdomain interactions within a single heptamer unit in both ^Sym^CDTb and ^Asym^CDTb (Fig. 3).

### The heptamer core of ^Asym^CDTb and ^Sym^CDTb

We refer to the heptamer domains (HD1, HD2, HD3) of aCDTb as the heptamer core because these regions of aCDTb retain a folds similar to that observed for other toxins in this class with their sequences aligning with up to ~20% identity. The first heptamerization domain (HD1) belongs to the clostridial calcium binding domain family (53). HD1 in both ^Sym^CDTb and ^Asym^CDTb features two proximal Ca^2+^ binding sites (Fig. 6A) that are highly conserved in this toxin family (54, 55), and the presence of Ca^2+^ was confirmed here for aCDTb using inductively coupled plasma mass spectrometry (ICP-MS). These Ca^2+^-binding sites do play a structural role in anthrax toxin (56, 57) and extracellular calcium is required for several steps in the intoxication of anthrax and iota toxin in cell-based assays (58, 59), so these results were not too surprising here for aCDTb. HD1 is followed by the β-barrel domain (βBD), and it is this domain that establishes the ~105 Å long β-barrel structure that is observed in what is termed here the “β-barrel heptamer unit”. Specifically, two strands from seven aCDTa subunits are elongated into 70 residue long double stranded anti-parallel β-sheet that together form this striking β-barrel fold. At the tip of the β-barrel, there are several hydrophobic residues that are partially protected from solvent via insertion into a cavity that presumably stabilizes ^Asym^CDTb prior to CDTa binding and/or insertion into the lipid membrane of host cells (Fig. 7A). While the β-barrel structure observed here for ^Asym^CDTb is reminiscent of the pore-forming component of the PA in the anthrax toxin, it is important to emphasize that in the case of ^Asym^CDTb, it does not require a lipid bilayer or presence of detergents to form. In the “non-β-barrel heptamer”, the βBD has a drastically different structure, as it retains a 4-stranded anti-parallel β-sheet that packs against HD2 and HD3, and this β-sheet structure is interrupted by a long loop that packs in between the third heptamerization domain (HD3) and the first receptor binding domain (RBD1).

The second heptamerization domain of aCDTb (HD2) has two anti-parallel β-strands followed by a 40-residue long loop, a short α-helix and a third β-strand, which completes a 3-stranded antiparallel β-sheet in both the non-β-barrel/β-barrel heptamers (Fig. S8). However, because HD2 packs into βBD of β-barrel heptamer, a rigid body type shifts in all three β-strands and the short helix of HD2 are observed that essentially “clamps down” on two strands of βBD to provide the unique packing of the “β-barrel forming” heptamer (Fig. S8). Whereas, in the non-β-barrel heptamer, this same HD2 domain is more “open” as it packs against all four strands of the β-sheet in βBD. Remarkably, this subtle difference in structure of HD2 is sufficient to reorient a key φ-gate residue, Phe-455, which is functionally important for transporting CDTa through the CDTb pore (37, 48). Thus, 7 phenylalanine residues in the φ-gate of β-barrel unit form a 3 Å orifice in comparison to the non-β-barrel units in which the pore diameter comprising these same phenylalanine residues is 12.5 Å (Figs. 7B, 7C). The final components of the heptamer core comprise a short 3 kDa turn-helix linker domain (LD1) and the third heptamerization domain (HD3). HD3 contains a four-strand β-sheet flanked by an extended loop region and two α-helices, and like the other two heptamerization domains, HD3 contributes to the large CDTa binding cavity just prior to the φ-gate (Fig. S9). For HD3, there are no significant structural differences between non-β-barrel and β-barrel heptamer units (r.m.s.d. of 0.32Å) for ^Sym^CDTb and A^sym^CDTb, respectively.

### The receptor binding domains of ^Asym^CDTb and ^Sym^CDTb

The first receptor binding domain (RBD1) is unique to aCDTb and has no sequence or structural similarity with any corresponding domains from anthrax toxin or any other binary toxin of known structure. RBD1 is a ten-stranded β-sandwich having a fold most similar to what are termed bacterial carbohydrate-binding modules (CBMs; Fig. S10). A number of β-sandwich CBMs are reported to bind calcium (60), as was observed here for RBD1 in the X-ray crystal structure of ^Asym^CDTb (Fig. 7B). Interestingly, RBD1 is better resolved in the crystal structure since it is stabilized by crystal contacts, whereas evidence for Ca^2+^ occupancy in the same location in cryoEM density is somewhat obscure due to increased flexibility of these regions in solution. Secondly, when the sequence comprising the RBD1 domain was isolated (residues 616-744), it was completely unfolded as determined by extreme broadening and lack of chemical shift dispersion in a ^15^N-edited HSQC NMR experiment; however, upon the addition of Ca^2+^, the linewidth values narrowed and significant chemical shift dispersion appeared that is typical of a fully folded protein. Importantly, evaluation of chemical shift indices in the NMR data Illustrate that Ca^2+^-bound RBD1 folds into a secondary structure that is similar to that observed of the full-length construct (Fig. 6).

The second, C-terminal receptor binding domain (RBD2) is connected to RBD1 by a 12-residue linker (L2; residues 745-756). Little if any change in its fold is observed when RBD2 is compared among all of the heptamer units (r.m.s.d. of 0.35Å) or to a crystal structure of this same domain determined here in isolation (**Fig. S11).** When the ^Sym^CDTb and ^Asym^CDTb structures are compared, however, the location of RBD2 is very different. Specifically, in β-barrel heptamer of ^Asym^CDTb, this is because of the L2 linker position and because of the formation of the long β-barrel domain itself. Thus, RBD2 of the β-barrel forming heptamer is located much closer to the protein core as compared to its position in the other heptamer of ^Asym^CDTb or to either heptamer of ^Sym^CDTb. This shift is combined with rotation of the entire donut-like structure as L2 linker is repositioned from a linear to angled orientation (Fig.4).

### The biological importance of receptor binding domain 2 (RBD2)

Importantly, RBD2 was found to be essential for promoting the di-heptamer assembly in both ^Sym^CDTb and ^Asym^CDTb since a ΔRBD2-CDTb construct, which lacks this C-terminal domain (residues 212-751), was found here to exist as a 7-subunit heptamer and not as a 14-subunit di-heptamer, as determined via SEC-MALS (data not shown). ΔRBD2-CDTb also had significantly reduced toxicity in Vero cell killing assays, even at concentrations greater than 10 µM (Fig. 1). Likewise, the isolated RBD2 construct (residues 757-876) behaved as a dominant negative and protected against a 500 pM dose of the binary toxin in *Vero* cell killing assays (IC_50_= 20nM ± 10 nM), (Fig. 1). Additionally, when challenging intact binary toxin with high concentrations of CDTa and ΔRBD2-CDTb, there was no protection against killing from the intact binary toxin. Taken together, these data show that the unique di-heptamer assembly involving RBD2 has an important role in the binary toxin’s biological activity and represents a domain in aCDTb worthwhile to target via structure-based drug design approaches.

### Possible biological role of the aCDTb di-heptamer

The core domain structures of each heptamer unit can be predicted based on similarities in sequence to other binary toxins (i.e. HD1/βBD/HD2/HD3) including 41% sequence identity to the corresponding region of anthrax toxin. Importantly, however, RBD1, has no sequence similarity to the corresponding receptor binding domain of anthrax toxin, and the anthrax toxin lacks the RBD2 domain altogether. Based on the pre-entry crystal structure of anthrax toxin, it was anticipated that aCDTb would also be heptameric in structure, particularly since the activity of CDTa:CDTb ratio was optimal at a 1:7 stoichiometry. Thus, the discovery of not one but two unique di-heptamer structures for aCDTb was very surprising. Demonstrating the exact nature of the evolutionary advantage conveyed by heptamer dimerization is beyond the scope of this study, but several possibilities can be outlined. The most intriguing of these is that pre-forming the β-barrel heptamer unit conformation, as found in ^Asym^CDTb, may facilitate toxin activity by accelerating its insertion into the membrane and that this process may be facilitated by binding to the host cell receptor. The second heptamer unit in this scenario, the non-β-barrel heptamer, could play the role of a “cap”, protecting and stabilizing the pore-forming heptamer of ^Asym^CDTb, as may be needed to increase its half-life *in vivo*. Another surprising discovery is that a second novel di-heptamer structure was discovered, ^Sym^CDTb, and this structure also needs to be considered in mechanistic terms. The essential stabilizing element of this structure is again the central donut shaped RBD2 tetradecamer. While the biological role of the symmetric structure is not fully understood, it may facilitate binding to CDTa, host cell receptor and/or host membrane binding. It needs to be pointed out that all of the biophysical data (SEC-MALS, SAXS, AUC) indicated that monomeric aCDTb (75 kDa) is still the major species (95±2%) in solutions of aCDTa, with the novel fourteen-subunit oligomer (1.0 MDa) detected at lower levels (<4-6%; Fig. 2). This result suggests that monomeric and di-heptameric forms of aCDTb are in a dynamic equilibrium. With this in mind, interconversion between the ^Sym^CDTb and ^Asym^CDTb may possibly follow two pathways, one direct (^Sym^CDTb ↔ ^Asym^CDTb), which was modeled here via normal mode analyses calculations (**Figs. S12-S14**), and another via dissociation into the monomeric form (^Sym^CDTb ↔ monomer ↔ ^Asym^CDTb). There was no evidence for a heptameric state of aCDTb, in any of the sizing studies used here, so aCDTb is unique when compared to other members of the binary toxin family.

### Summary

The structures of ^Sym^CDTb and ^Asym^CDTb reported here show several novel and unexpected features that provide routes towards rational drug design aimed at inhibiting the *C. difficile* binary toxin. Such strategies are much needed since there is nothing approved by the FDA to target CDT (12). Therefore, to address this unmet medical need, identifying vulnerabilities in CDT are needed for developing novel therapeutic strategies to target CDT. With these two novel structures in hand (Fig. 3), the unexpected dimerization of two heptameric assemblies via extensive interactions in the RBD2 domain make this a particularly promising region to target. Likewise, blocking other regions (i.e. RBD1, Ca^2+^-binding domains, other domains) could also turn out to be lethal for the toxin. Nonetheless, the structures reported here CDTb indicates that *C. difficile* binary toxin must use a different mechanism for delivering its enzyme payload, CDTa, into target cells versus other heptameric toxins such as the PA from the anthrax toxin, so unique strategies to inhibit this toxin’s lethal activity could benefit from the structural information reported here.

A second important component of this work is the synergistic approach to structural characterization of aCDTb. Several structural and biophysical methods were employed that provided a multi-faceted examination of the problem. Cryoelectron microscopy is the nexus of this work as it provided the initial discovery of the aCDTb di-heptamer in two different conformations even for a low percentage of the protein (4-6%). Knowledge of these structural assemblies from cryoEM then allowed for resolving phasing issues in the crystal structure determination, which provided feedback regarding novel Ca^2+^-binding site in the RBD1 domain. Furthermore, NMR techniques were employed to indicate that RBD1 Ca^2+^-binding is likely important for its stability. Additionally, the ability to detect multiple conformations of the aCDTb in solution by cryoEM enhanced the analysis of the SAXS data, which originally provided impetus for considering higher oligomerization states as radius of gyration data was inconsistent with a heptamer models based on homology. Even initial models provided an improved fit once conformational heterogeneity was included in the analysis. SAXS data also confirmed that the dual conformation is present in solution and is not an artifact of freezing procedure employed in preparing EM samples. Lastly, other biophysical techniques (SEC-MALS and AUC) for characterizing size distributions in solution indicated that a significant amount of monomeric protein is present in the solutions used for these structural studies of aCDTb (>90%). Noteworthy, the structural methods employed here are all insensitive to this for different reasons. CryoEM analysis is based on picking particles in the micrographs and is thus dominated by larger clearly discernible megadalton size di-heptamer. In SAXS, larger particles dominate scattering intensity with the detection of smaller monomers being negligible. X-ray crystallography resolves structures that crystallize. In this case, it is remarkable that a conformation that probably represents no more than 2% of the protein particles is the one that crystallized, upending the traditional notion that highly concentrating monodisperse protein is a pre-requisite of a successful crystal structure determination. Lastly, without knowledge of the protein size distribution (>90% monomer) under the conditions of this structure-determination work, a starkly different picture may have arisen for how to describe the transition between the two di-heptamer conformations (**Figs. S12-S14;** ^Sym^CDTb ↔ ^Asym^CDTb), but these data forced consideration that the conversion between ^Asym^CDTb and ^Sym^CDTb could be mediated by an oligomer assembly/disassembly mechanism from the monomeric form (i.e. ^Sym^CDTb ↔ monomer ↔ ^Asym^CDTb). Furthermore, since monomeric pro-CDTb is not toxic, it opens up a new therapeutic possibility, as it suggests that fully assembled aCDTb is in active equilibrium, which may be potentially shifted by small molecule inhibitor(s) or biologics. Thus, capitalizing on such a multi-faceted approach to molecular characterization, it was shown that the resulting picture from the multiple methods is more than the sum of its parts, particularly for large macromolecular assemblies such as aCDTb (> 1 MDa). In summary, the individual structural methods (cryoEM, X-ray crystallography, NMR) provide phenomenal insights on their own, but they become even more powerful when used together and when combined with other biophysical techniques.

## MATERIALS AND METHODS

### Protein expression and purification

Active CDTb (aCDTb) was expressed and purified as described in (12). Briefly, full length pro-CDTb was expressed in insect cells-baculovirus system and purified using affinity chromatography. To obtain the active protein, the N-terminal activation domain was proteolytically removed using chymotrypsin and purified by size exclusion chromatography. Full-length CDTa and several truncated constructs of CDTb (RBD1, RBD2, and ΔRBD2-CDTb) were overexpressed in *E. Coli* and purified to homogeneity by combination of affinity and size exclusion chromatography methods as described in the Supporting Information (SI).

### Vero cell activity assay

Briefly, *Vero* cells incubated in presence of binary toxin were quantified for F-actin using fluorescently labeled phalloidin to determine toxicity. Further details are provided in SI.

### Cryoelectron microscopy

Purified aCDTb was placed on holey gold grids with an additional thin layer of carbon on top, blotted and flash frozen in liquid ethane using FEI Vitrobot IV. Grids were inspected and electron micrographs collected on FEI Titan Krios at 300V equipped with Gatan K2 Summit direct electron detector. Multiple iterative rounds of 2D/3D classification resulted in identification of two distinct protein conformations for which the density maps were refined with Bayesian particle polishing and CTF refinement with Relion (61) to 3.14 Å and 2.84 Å resolution. Further details are provided in SI.

### X-ray crystallography

Crystallization conditions were found for all the structures via sparse matrix robotic screening. Standard techniques of cryoprotection were used and experimental diffraction data was collected at SSRL. Structures were solved by molecular replacement using PHASER (62) and refined with phenix.refine (63). Further details are provided in SI.

### Nuclear magnetic resonance

A 2D ^15^N-edited HSQC of 0.5 mM RBD1 in 15mM HEPES (pH 7.0), 150 mM NaCl, 10% D_2_O was collected at 950 MHz, 25°C. Minimal residues appeared with high noise. 2.3 mM Ca^2+^ was added and the ^15^N-edited HSQC was collected under the same conditions. The spectrum cleared and the number of residues noticeably increased. The Ca^2+^ concertation was raised to 6 mM and the spectrum improved further with no additional changes at higher Ca^2+^ concentrations (>12 mM).

### Biophysical techniques

Biophysical characterization of the activated CDTb included small angle X-ray scattering, size exclusion chromatography with multi angle light scattering analysis (SEC-MALS) and analytical ultracentrifugation (AUC). Experimental details for these techniques are described in the SI.

## DATA AVAILABILITY

Cryo-EM density maps have been deposited in the Electron Microscopy Data Bank (EMDB) under accession numbers EMD-20926 for ^Asym^CDTb, and EMD-20927 for ^Sym^CDTb. Model coordinates have been deposited in the Protein Data Bank (PDB) under accession numbers 6UWR for ^Asym^CDTb, 6UWT for ^Sym^CDTb, 6UWI for crystal structure of the full length CDTb in ^Asym^CDTb conformation, and 6UWO for crystal structure of the RBD2 domain of CDTb. All other data are available from the corresponding authors upon request.

## ACKNOWLEDGMENTS

Funding from the Center for Biomolecular Therapeutics (CBT; to DJW), Maryland Center for Advanced Molecular Analyses (M-CAMA; to DJW, EP) and the at the University of Maryland School of Medicine are acknowledged. A.d.G acknowledges funding from the NIH (grant no. R35GM133598) and CUNY. Funding to support the ICP-MS analyses was from the National Science Foundation (NSF CHE 1708732). Use of the Stanford Synchrotron Radiation Lightsource, SLAC National Accelerator Laboratory, is supported by the U.S. Department of Energy, Office of Science, Office of Basic Energy Sciences under Contract No. DE-AC02-76SF00515. The SSRL Structural Molecular Biology Program is supported by the DOE Office of Biological and Environmental Research, and by the National Institutes of Health, National Institute of General Medical Sciences (P41GM103393). The contents of this publication are solely the responsibility of the authors and do not necessarily represent the official views of NIGMS or NIH.

