## Supplementary Information for "Structure of the cell-binding component of the *Clostridium difficile* binary toxin reveals a novel macromolecular assembly"

**Supplementary Information (Xu *et al.*, 2019)**

*Vero Cell Toxicity Assay*. Vero cells (ATCC CCL-81) were obtained from American Type Culture Collection and cultured in EMEM (Corning Cellgro 10-009-CV) supplemented with 10% FBS (i.e. complete media). Cells were seeded at a density of 2000 cells/well in 200 µl complete media in 96-well format (Corning #3904) and incubated overnight in a humidified incubator with 5% $CO_2$ at 37°C. For toxin dose-response, CDTa was combined with either aCDTb or activated ΔRBD2-CDTb to yield a 1:7 molar ratio stock solution of binary toxin (*i.e.* 1 µM CDTa with 7 µM aCDTb). Binary toxin stock was serially diluted in complete media and a final volume of 100 µl was added to the existing media in the 96-well plate at final concentrations up to 10 µM toxin. Cells were incubated with binary toxin for two hours in 5% $CO_2$ at 37°C and a dose-response curve was generated for each combination. For the RBD2 domain challenge, 500 pM binary toxin (500 pM CDTa: 3500 pM aCDTb) was challenged with the RBD2 domain at concentrations between 1 pM -100 µM and incubated for 1 hour. After indicated incubation times, plates were centrifuged at 1000 RPM for 1 minute. Media was removed and cells were fixed by incubating in 3.7% Formaldehyde in 1x PBS for 15 minutes at room temperature, followed by a wash and incubation with 1% Triton-X-100 in 1xPBS for 5 minutes at room temperature to permeabilize the cells. Cells were washed and incubated in the dark for 1 hour at room temperature with 0.016 µM Alexa Fluor 488 Phalloidin in 1x PBS + 1% BSA, washed and incubated with 1xPBS for imaging. Fluorescence of Alexa Fluor 488 Phalloidin was read in well-scanning mode using the BMG PHERAstar FS with excitation 495 nm, emission 518 nm. Normalized Fluorescence was plotted against the concentration of CDTa. Data was graphed in GraphPad Prism 6.0 with $TC_{50}$ and $IC_{50}$ values reported from standard curve fitting and with errors reported from analyses of multiple experiments ($n \geq 3$; as indicated).

*EM sample vitrification and data acquisition*. Purified aCDTb protein sample was adjusted to 55 μM with 50 mM HEPES pH 7.5 and 150 mM NaCl. 3-μL aliquot was blotted onto a glow-discharged QUANTIFOIL® UltrAuFoil R 1.2/1.3 grid with a thin carbon layer on top (QUANTIFOIL, GMBH) using a FEI Vitrobot IV with a blotting time of 1s, waiting time of 0s, 100% humidity, and a blotting force of 4. The grid has been plasma cleaned using a Fischione M1070 NanoClean before applying the sample. The sample was flash frozen in liquid ethane. Frozen grids were stored in liquid nitrogen before use.

aCDTb containing grid was loaded onto a 300 keV Titan Krios electron microscope (FEI, Eindhoven), and the micrographs were collected automatically with a Gatan K2 Summit direct electron detector with a pixel size of 5 microns and a calibrated magnification of 47170x. A total of 1204 micrographs were collected with pixel size at the object scale of 1.06 Å and a defocus range of 0.5-6.7 μm. 40 frames per micrograph were collected, with a dose rate of 8 electrons, which translates to total dose of 56.9 electrons per $Å^2$. Detailed data collection and processing parameters are shown in Table S1.

*Low resolution initial model generation*. Initial micrographs were collected with gold grids without thin carbon layer. All particles preferred top down orientation, revealing only the top view of the protein complex. The micrographs were motion corrected with MotionCor2 (1). Contrast Transfer Function (CTF) estimation was done with a GPUaccelerated CTF estimation software (GCTF v1.18) using all frames (2). 2D classification confirmed that only the top view classes are present from the collected data. To resolve this issue, a tilting strategy was applied. The images were collected at 40° angle, the micrographs were gain and motion corrected the same way as the original data (3). CTF estimation with GCTF was done to accurately assess the defocus values for the tilted micrographs. 2D classification revealed multiple classes of particles with different orientations as a result of the tilting scheme. *ab initio* initial model generation with only the tilted particles was performed with cryoSPARC v1 (4) without success. Particles form the top view classes and the tilted particles were then pooled together to form a set of 24585 particles (ratio was: top view 1: tilted 9). This allowed the generation of the first sub nanometer resolution model of the CDTb complex (~6Å).

*High resolution density maps generation*. We later tested and confirmed that by using gold grid with a thin carbon layer can substantially improve the distribution of orientations of the particles on the grids. We collected the new dataset (1204 micrographs) without tilting. Motion correction and CTF estimation were done in an identical manner as previously described. Image processing was carried out with Relion 2.0 and cryoSPARC v1 (4-6). In Relion, 845 particles were manually selected, and extracted as reference for automated particle picking. A total of 68972 particles were picked with a box size of 400 pixels. The particles were re-scaled to 150 pixels for subsequent 2D classifications. The number of 2D classes was initially set to 60, with diameter of the mask set at 340 Å. Limit resolution E-step was initially set to 30 Å, but gradually reduced to 15 Å after every 3 – 7 iterations. Setting a resolution limit for the expectation step (i.e. E-step) in the beginning prevents overfitting of the data, which showed increased alignment speed and improved classification results by yielding more diverse sets of views in 2D or more diverse set of conformations in 3D classification, while also increasing computational speed. A subset of 27 classes with 54040 particles were selected and a second round of 2D classification with 24 classes. The E-step was initially set to 20 Å but gradually reduced to 15 and then -1 over 8 iterations. A total of 17 classes with 41735 particles were selected and re-extracted with the original box size and uploaded to cryoSPARC for initial model generation and 3D classifications. In cryoSPARC, the uploaded particles were used for *ab initio* model generation with 10 classes. The classes generated revealed two distinguishable initial models including a compacted form ($^{Asym}$CDTb) and a more extended form ($^{Sym}$CDTb), with 11448 particles and 21242 particles respectively. Homogeneous refinement in cryoSPARC using the two initial models yielded two high resolution density maps, at 3.5 Å and 4.1 Å resolution. 3D refinements of the two structures were also carried out in Relion with initial low-pass filter at 50 Å, which yielded the density maps at 3.6 Å and 4.2 Å. The maps obtained from the two applications share similar density volumes. The maps obtained appear to be following C7 symmetry. C7 symmetry refinements of the two structures were carried out in cryoSPARC and yielded density maps with improved resolutions, at 3.1 and 3.3 Å resolution. Resolutions were estimated based on the Fourier Shell Correlation (FSC) with 0.143 as cutoff.

To improve the resolution of the density maps further, particles from micrographs with higher defocus values were removed. A set of defocus value cutoff were tested, and a defocus value cutoff of 2 microns was found to be optimal. This reduced the total number of particles in each of the two classes to 10931 particles and 17406 particles for the $^{Asym}$CDTb and $^{Sym}$CDTb conformations, respectively. Homogenous refinement using the filtered set of particles yielded a final resolution of 3.0 Å for $^{Asym}$CDTb. The density map for $^{Sym}$CDTb did not show discernable improved resolution.

*CTF refinements and particle polishing*

Next, we explored the CTF and Beamtilt refinement and Bayesian polishing steps newly implemented in Relion 3.0 (7). We first motion corrected the micrographs with Relion's built-in MotionCor2. We grouped 3 frames together for beam-induced shifts calculation for better initial results. CTF estimation was carried out as previously described using GCTF. 2D classes from previous 2D manual picking were used as the references for a new auto-picking step with reduced threshold. A total of 73230 particles were picked and extracted with re-scaled box size of 150 Å. 2D classifications were done as previously described. 2D averages included top views, side views, and tilted views of the particles. 13 classes containing 62786 particles were selected and re-classified into 25 classes. 61711 particles from 21 classes were selected for 3D classification using the previously obtained $^{Asym}$CDTb density map as the reference. The particles were re-extracted to the original box size (400 Å) prior to 3D classifications. 3D classifications with 10 classes revealed a single $^{Asym}$CDTb from class, and multiple classes representing the $^{Sym}$CDTb form. The $^{Sym}$CDTb form classes were selected to undergo a 2$^{nd}$ round of 3D classification with 6 classes. Two classes containing 19944 particles similar to each other showed good agreement with the previously obtained density map corresponding to the $^{Sym}$CDTb form, and the particles were pooled for subsequent refinements. 11122 particles from the $^{Asym}$CDTb conformation 3D classification were used for 3D auto-refinement in Relion using the 3 Å density map obtained previously as reference, which resulted in a 3.2 Å density map post sharpening. CTF and beamtilt refinement was applied to the particles, followed by Bayesian polishing. During the polishing step, 5000 particles were used in the training mode to generate optimized parameters. Polished particles were refined again to obtain a 2.9 Å resolution map post sharpening. Two iterations of CTF refinements were applied to the particles and the resolution of the density map converged at 2.8 Å. Identical processes were applied to the $^{Sym}$CDTb conformation particles with the previously obtained 3.3 Å map as reference. The resolution of the density map improved from 3.4 Å and converged at 3.1 Å following three iterations of CTF refinements and particle polishing. Example of micrographs with top and side views, 2D and 3D averages, FSC curve of the final density maps, and 3D reconstruction maps of the two conformations are shown in Figure 1S. Local resolution calculated from ResMap are also displayed on the sharpened maps (8). All 3D maps are visualized by UCSF Chimera (9, 10).

*Size determination by Size Exclusion Chromatography Multi Angle Light Scattering (SEC-MALS)*. For SEC-MALS, a UHPLC system (Vanquish Flex, Thermo Fisher) was coupled to MALS (DAWN HELEOS-II, Wyatt) and Refractive Index (Optilab T-rEX, Wyatt) detectors. Separations were performed using a WTC-050N5 column (Wyatt) equilibrated in PBS, with a flow rate of 0.3 mL/min and sample injection volumes of 25 µL. Molar mass analysis was performed using the software ASTRA 7.1.3 (Wyatt) using refractive index as a concentration source. To determine the oligomerization state of the activated CDTb, it was analyzed using SEC-MALS. While the ~1MDa macromolecular assembly matching the size of aCDTb tetradecamer is clearly present, most of the protein materials remains in monomeric (~75kDa) form.

*Analytical ultracentrifugation studies (AUC)*. Sedimentation velocity measurements were carried out using a Beckman Coulter Optima XL-I analytical ultracentrifuge equipped with a 4-hole An-60Ti rotor. Measurements were performed on aCDTb protein prepared at 11 M in buffer containing 15 mM HEPES, pH 7, 150 mM NaCl at 20 deg C. The sample was centrifuged at 30,000 rpm at 20oC in a standard two-hole cell equipped with an epon charcoal-filled centerpiece. A total of 860 absorbance scans were acquired at 280 nm in continuous mode. The data were analyzed using DCDT+ to obtain a g(s*) distribution, which was subjected to fitting using a two species model to obtain estimates of species sedimentation coefficients and molecular weights. Values of protein partial specific volume and buffer density and viscosity were calculated using the program Sednterp (http://rasmb.org/sednterp/).

*Small angle X-ray scattering*. Solution scattering data for aCDTb were acquired using MOLMEX Ganesha instrument. Copper Kα incident radiation with wavelength of 1.542Å was produced by the Rigaku MicroMax 007HF rotating anode generator, monochromated and collimated via 2 pinholes equipped with scatter-less slits. Pilatus 300K detector was used to register scattered radiation with the transmitted intensity monitored via a pin diode. Samples were kept at 25 ºC and exposed to incident X-ray radiation for 12 sequential 15-min frames. Pixel outliers due to background radiation were removed and the 2D data corrected for detector sensitivity and solid angle projection per pixel. The data were converted to one-dimensional scattering intensity curves, frame-averaged and buffer-subtracted. Data sets acquired at sample-detector geometries of 1035 mm and 355 mm were merged to extend the angular resolution range. In addition to the stock concentration of 2 mg/mL, scattering data were acquired for two- and four-fold diluted samples in order to investigate concentration dependence. As no such dependence was found by the Guinier and P(r) analyses for the two lowest concentrations tested, scattering curves at 1 mg/mL protein concentration were used for further work. Guinier fit of the lowest-angle scattering data resulted in he fitted radius of gyration of 86±2 Å and zero-angle scattering intensity corresponding to the protein molecular mass of 1.0±0.2 MDa by reference to scattering signal of water measured at identical conditions after subtraction of the empty capillary scattering. Estimation of the total volume of the scattering particle via Porod invariant resulted in the excluded volume of 1.3 MÅ$^3$ corresponding to protein molecular mass of 1.1 MDa. Fourier transform of the scattering data yielded gyration radius of 85±1 Å, consistent with the results of the Fourier analysis and indicating sample monodispersity. Inter-atomic distance probability distribution fitted to the I(q) data, shown in the paper along with the corresponding I(q) fit, indicates maximum particle dimension of 270 Å and suggests dumbbell shape of the scattering particle.

*aCDTb crystallization and X-ray data collection*. Robotic sparse-matrix screening was performed to obtain initial crystallization conditions for aCDTb. These were further optimized to obtain X-ray quality crystals. Large crystals grow over several days in a range of conditions that include 22-27% PEG1500 and 0.1M MIB buffer at pH 6-8. Crystals were cryoprotected by briefly incubating them in 40% PEG1500 and 0.1M MIB buffer at pH 7, flash-cooled in liquid nitrogen and stored for data collection. X-ray data were collected at SSRL beamline 12-2, and processed using XDS and AIMLESS. Most crystals diffracted to 5-7Å and a large numbers of datasets had to be collected to find those that occasionally diffracted to better than 4Å. Since crystals were fairly large (up to 300μm), in all cases multiple datasets were collected from non-overlapping sections of a crystal that were then merged in order to improve data quality. Ultimately, we were able to obtain a dataset from a single crystal with the resolution limit as high as 3.7Å. Data processing statistics are shown in Table S2.

*Structure determination*. Crystal structure of aCDTb was solved by molecular replacement using model of the aCDTb obtained by cryoEM. Initially, space group was determined as C2 based on symmetry of the diffraction patterns alone. However, while molecular replacement using PHASER did produce a solution characterized by reasonably high log-likelihood gain, it was rejected due to large number of self-clashes. The issue was resolved by reducing the symmetry to triclinic lattice. To solve the structure, cryoEM model was split into four structural domains: core heptamers of both $^{Sym}$CDTb and $^{Asym}$CDTb conformations, β-barrel, and RBD2 tetradecamer. These were combined sequentially by molecular replacement, which provided additional validation as individual domains packed in the way that was consistent with cryoEM model. As expected, only one of the two conformations are present in the crystal – that of the $^{Asym}$CDTb. Initial refinement revealed several structural differences between crystal structure and cryoEM model. In particular, a significant peak in the difference density was located in all copies of the RBD1 domain that was consistent with additional calcium atom. ICP-MS analysis shows that calcium is the only metal present in active CDTb, hence calcium was placed in the structure model. CryoEM density is consistent with this additional metal binding site and it is thus not an artifact of crystallization. It was, however, overlooked while building the cryoEM model, in part because presence of the metal ion is more obvious in difference density, and also because of increased disorder in RBD1 domain. In crystal structure, RBD1 domains provide the (only) crystal contact and therefore appear more ordered. Furthermore, the core heptamer of the β-barrel conformation appears to be slightly rotated in the crystal structure (apparently due to optimization of crystal contacts), and electron density suggests that the LD2 linker packs in a compact conformation, generating a "linear" arrangement of the RBD1/RBD2 domains similar to one found in non-β-barrel conformation (see Fig. S3). Additional unexplained electron density was also found in the gap between RBD1 and RBD2 domains. Detailed analysis revealed that this additional density is consistent with the CDTb tetradecamer that is flipped 180 degrees. Slight rotation of the core heptamer results in perfect overlap of RBD1 domain in these alternative orientations, which in turn preserves crystal contacts. This also renders the crystal structure with perfect two-fold symmetry (relative occupancy of the two orientations was selected in series of refinements with fixed occupancies by choosing the occupancy value that results in the lowest $R_{free}$), which explains why initial analysis of raw diffraction data yielded higher symmetry of predicted space group. The crystal structure of $^{Asym}$CDTb provides validation of the cryoEM model. It does, however, only identify one of the two conformations, with the $^{Sym}$CDTb di-heptamer being incompatible with the crystal lattice. Rotation of the core heptamer relative to the RBD2 tetradecamer demonstrates conformational flexibility of the protein, which is only partially captured in the current cryoEM model.

*RBD2 domain crystallization and X-ray data collection*. Robotic sparse-matrix screening was performed to obtain initial crystallization conditions for isolated RBD2. These were further optimized to obtain X-ray quality crystals. Large crystals grow over 1-2 weeks with mother liquor consisting of 0.2 M sodium chloride, 1.0 M sodium citrate and 0.1 M Tris pH 7.0. Crystals were cryoprotected by transferring them into 1.6 M sodium citrate, flash-cooled in liquid nitrogen and stored for data collection. X-ray data were collected at SSRL beamline 12-2, and processed using MOSFLM and AIMLESS. Data processing statistics are shown in Table S2.

*RBD2 structure determination*. Straightforward molecular replacement using PHASER with the RBD2 domain extracted from the cryoEM structure as a model produced good quality electron density maps that required only minor manual rebuilding and refinement. Refinement statistics are shown in Table S2.

NMR spectroscopy. NMR data was collected at 25°C on Bruker Avance III 600 MHz and 950 MHz spectrometers, both equipped with TCI cryogenic probes. The NMR sample contained 0.5 mM RBD1 protein, 15 mM HEPES buffer (pH 7.0), and 150 mM NaCl in 90% $H_2O$ and 10% $D_2O$ with data collected at 25 °C. Addition of 6 mM $Ca^{2+}$ was found to dramatically improve the signal to noise ratio and the number of disperse observable residues. For resonance assignments of $Ca^{2+}$-loaded RBD1, triple resonance heteronuclear 3D HNCA, HN(CO)CA, HNCACB with CB resonances optimized, CACB(CO)NH, CC(CO)NH, HNCO and HN(CA)CO 3D experiments were collected. The data was processed with NMRPipe (11) and analyzed using CcpNmr (12) software. The secondary structure was predicted using the CSI 2.0 web server (13).

Normal mode analysis calculations. Normal mode analysis calculations were undertaken to identify possible global motions that may contribute to the transition between these two experimentally observed $^{Sym}$CDTb and $^{Asym}$CDTb conformations. Two normal modes are observed to make substantial contributions to the confrontational changes from which a possible pathway between the two conformations of the CDTb heptamer is identified. As shown in Fig. S12, the first step of the conformational transition involves rearrangement of several domains including heptamerization domain II, receptor binding domain I, linker domain II and receptor binding domain II. Rotations of these domains allows the beta-barrel domain to access a wider range of conformations allowing for formation of the beta-barrel. Receptor heptamerization domain II in conjunction with mediating waters is hypothesized to allow for malleability of the contacts between beta-barrel domains from different monomers thereby promoting the formation of the beta-barrel, thereby leading to the ability of the protein to deliver toxic enzymes into cytosol when inserted into the endosomal membrane. Likewise, examination of each individual domains provides insights regarding the mechanism of action for aCDTb, which includes calcium binding (HD1, RBD1), formation of a pore-forming β-barrel domain that modulates the position of 7 phenylalanine residues in the φ-gate (βBD, HD2), and importantly, the domains involved in di-heptamer assembly as well as host cell receptor binding (RBD1, RBD2). Normal mode analysis (NMA) calculations and following overlap analyses were conducted to explore possible confrontational changes between the two CDTb conformations for both heptamer and 14-mer structures. NMA for the heptamer was performed using elNemo sever (14) and more expensive NMA for 14-mer was done with WEBnm@ sever (15). After NMA calculations, overlap analyses were performed to check important normal modes that are contributing significantly to the related conformational changes.

Figure S13 shows the overlap and cumulative overlap plots for normal models that were calculated for heptamer of CDTb. Two normal modes 34 and 30 has the most overlaps with the conformational change vectors and are predicted to be highly related to the conformation change between two conformations. Normal mode 34 is related to a breathing like mode that beta barrel is vibrating up and down and the crown is contracting and expending. Normal mode 30 is seen to be rotations of different components in the crown part and related to the rearrangement between different heptamerization domains. The RMSDs related to these two normal modes were mapped onto structures and are shown in Figure S14. It can be seen directly that large RMSD changes are seen for both crown and barrel parts for mode 34, while only seen for crown part for mode 30. Similar behavior is seen for 14mer case. Normal mode 43 and 56 are found to overlap the most with conformational change vectors. And similar crown breathing and beta barrel up and down vibration is seen for mode 56 and similar component rearrangements is seen for the crown part for mode 43. Thus, for both heptamer and 14mer case, similar types of vibration modes are found to be the most related to the conformation change between the two conformations.

TABLES

Table S1. Details of cryoEM data collection.

| | SymCDTb | AsymCDTb |
|---|---|---|
| **EMDB ID** | EMD-20927 | EMD-20926 |
| **PDB ID** | 6UWT | 6UWR |
| **Microscope** | FEI Titan Krios | |
| **Voltage (kV)** | 300 | |
| **Spherical aberration (mm)** | 2.7 | |
| **Amplitude Contrast** | 0.07 | |
| **Camera** | Gatan K2 Summit | |
| **Defocus range all data (µm)** | 0.5-6.7 | |
| **Defocus mean ± std all data (µm)** | 1.4 ± 0.6 | |
| **Nominal defocus range used (µm)** | 0.5 - 2.5 | 0.6 - 2.5 |
| **Defocus mean ± std used (µm)** | 1.2 ± 0.3 | 1.3 ± 0.3 |
| **Exposure time (s)** | 8 | |
| **Dose rate ($e^{-}$ $pixel^{-1}$ $s^{-1}$)** | 8 | |
| **Total dose ($e^{-}$ $Å^{-2}$)** | 56.9 | |
| **Pixel size (Å)** | 1.06 | |
| **Number of micrographs (total)** | 1204 | |
| **Number of micrographs (used)** | 1087 | |
| **Frames** | 40 | |
| **Total Particles Picked** | 73230 | |
| **Ice thickness (nm)** | 40-100 | |
| **Ice thickness average (nm)** | 70 | 70 |
| **Particles Used** | 19944 | 11122 |
| **Resolution Before Symmetry (Å)** | 3.5 | 3.9 |
| **Symmetry Imposed** | C7 | |
| **Resolution (Global) (Å)** | 3.1 | 2.8 |
| **Estimated b-factor ($Å^2$)** | -54 | -38 |

Table S2. X-ray data collection statistics and model quality. Values in parentheses correspond to the highest resolution shell.

| | $^{Asym}$CDTb | RBD2 |
|---|---|---|
| PDB ID | 6UWI | 6UWO |
| **Data collection** | | |
| Wavelength, Å | 0.979 | 0.979 |
| Space group | P1 | $P6_5$ |
| Unit cell parameters | a=190.5 Å, b=191.0 Å, c=192.3 Å<br>α=108.7°, β=94.5°, γ=108.0° | a=53.7 Å, b=53.7 Å, c=171.2 Å<br>α=90°, β=90°, γ=120° |
| Resolution range, Å | 39.8-3.70 (3.76-3.70) | 45.1-2.30 (2.38-2.30) |
| Multiplicity | 11.2 (9.9) | 17.7 (18.2) |
| Unique reflections | 237,933 (11,967) | 12588 (1228) |
| Completeness, % | 93.2 (94.8) | 99.8 (99.8) |
| $\langle I/\sigma_I \rangle$ | 4.2 (0.6) | 4.6 (1.0) |
| CC1/2 | 0.996 (0.341) | 0.989 (0.247) |
| **Refinement** | | |
| Resolution | 39.7-3.70 (3.742-3.700) | 44.9-2.30 (2.53-2.30) |
| No. of reflections | 234,146 (7,686) | 12,336 (2,965) |
| $R_{work}$ | 0.280 (0.399) | 0.240 (0.287) |
| $R_{free}$ | 0.296 (0.413) | 0.295 (0.345) |
| No. of atoms<br>Protein<br>Metal ions | <br>70,789<br>42 | <br>1,964 |
| Average B-factors, $Å^2$<br>Protein<br>Metal ions | <br>127.1<br>102.5 | <br>19.4 |

Figure S1. Screenshot of a typical micrograph with top and side views of CDTb complex on the gold grid with thin carbon film.

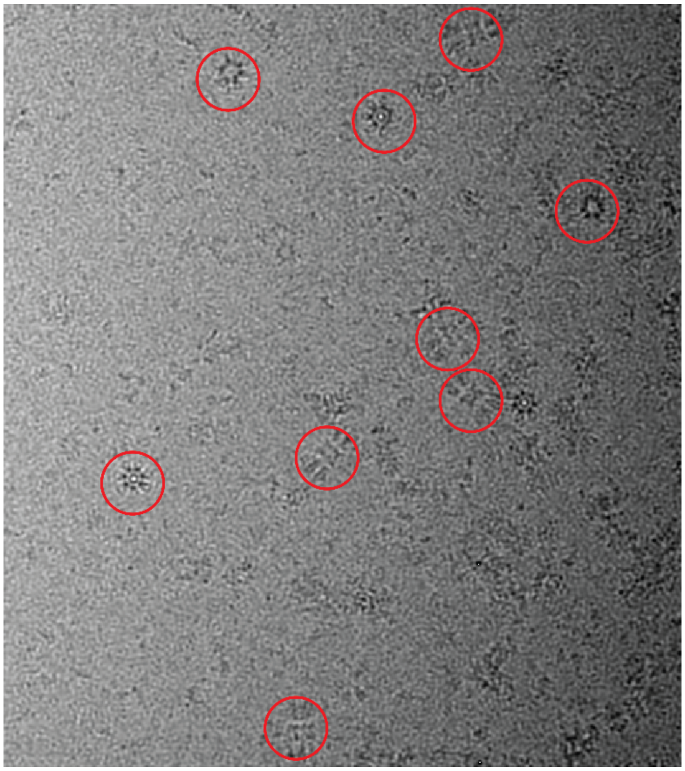

Figure S2. Workflow of obtaining density maps of the two conformations of aCDTb complex. a) Examples of 2D averages obtained from non-tilted (left) and tilted (middle) gold grid, and from gold grid with thin carbon film (right). Particles on gold grid with thin carbon film are found in top, side, and tilted orientations. b) 3D classes generated from RELION 3D classifications. Two classes representing the $^{Sym}$CDTb conformation were and one class of $^{Asym}$CDTb conformation were used for refinements of the two conformations. Sharpened final density maps of 3.1 and 2.8 Å are colored with local resolutions. c) Fourier Shell Correlation (FSC) curves for the final density maps ($^{Sym}$CDTb conformation on the left, and $^{Asym}$CDTb conformation on the right) are shown.

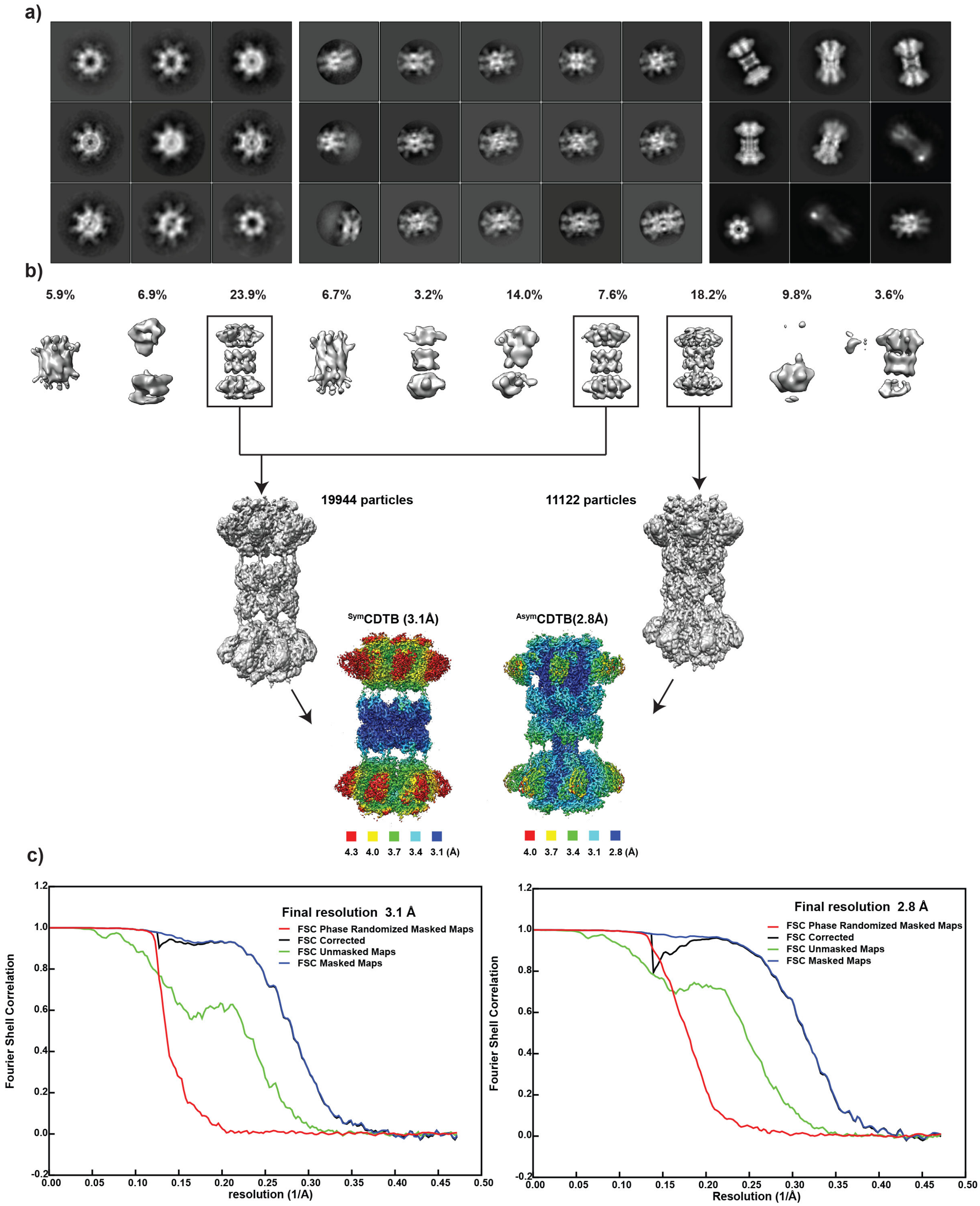
a)
b)
5.9%
6.9%
23.9%
6.7%
3.2%
14.0%
7.6%
18.2%
9.8%
3.6%
19944 particles
11122 particles
SymCDTB (3.1Å)
AsymCDTB(2.8Å)
4.3 4.0 3.7 3.4 3.1 (Å)
4.0 3.7 3.4 3.1 2.8 (Å)
c)
Final resolution 3.1 Å
FSC Phase Randomized Masked Maps
FSC Corrected
FSC Unmasked Maps
FSC Masked Maps
Fourier Shell Correlation
resolution (1/A)
Final resolution 2.8 Å
FSC Phase Randomized Masked Maps
FSC Corrected
FSC Unmasked Maps
FSC Masked Maps
Fourier Shell Correlation
Resolution (1/Å)

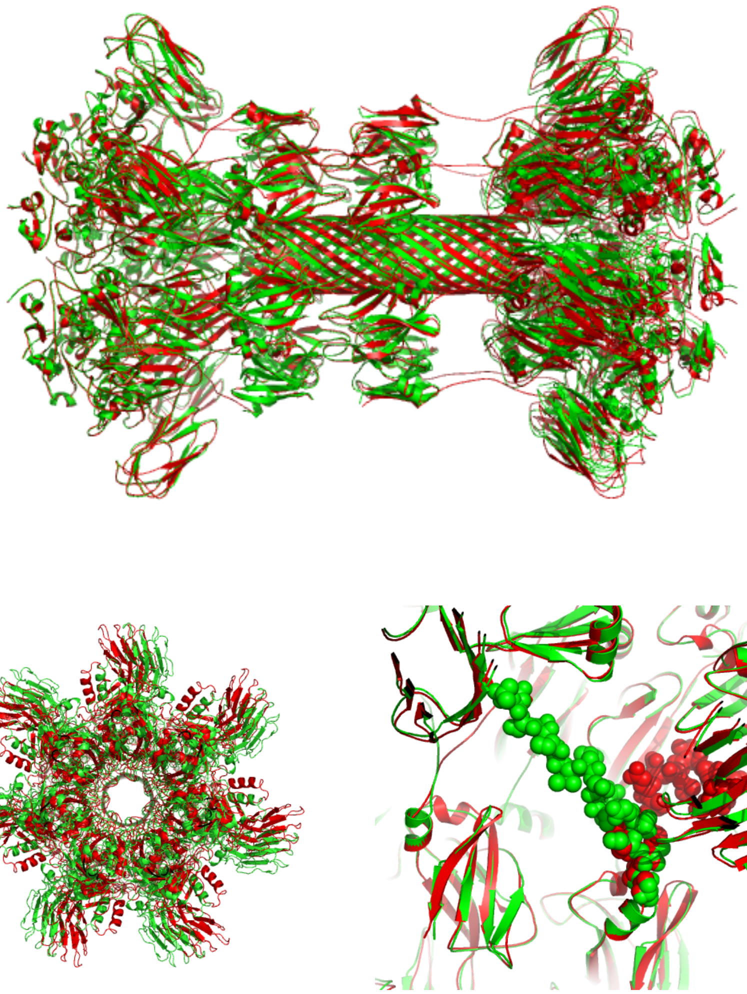

Figure S3. Comparison of the cryoEM (green) and crystal (red) structure demonstrates that the protein crystallizes in $^{Asym}$CDTb form. Bottom left shows a detectable relative rotation of the two heptamers. Bottom right panel shows re-arrangement of the linker connecting RBD1 and RBD2 producing “linear” arrangement of the RBD2 domains with respect to the heptamer core, similar to one observed in $^{Sym}$CDTb conformation.

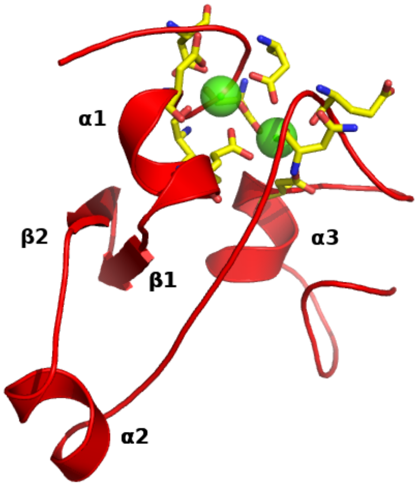


Figure S4. Heptamerization domain 1 structure.  Dual calcium binding site and metal ion ligands are shown.

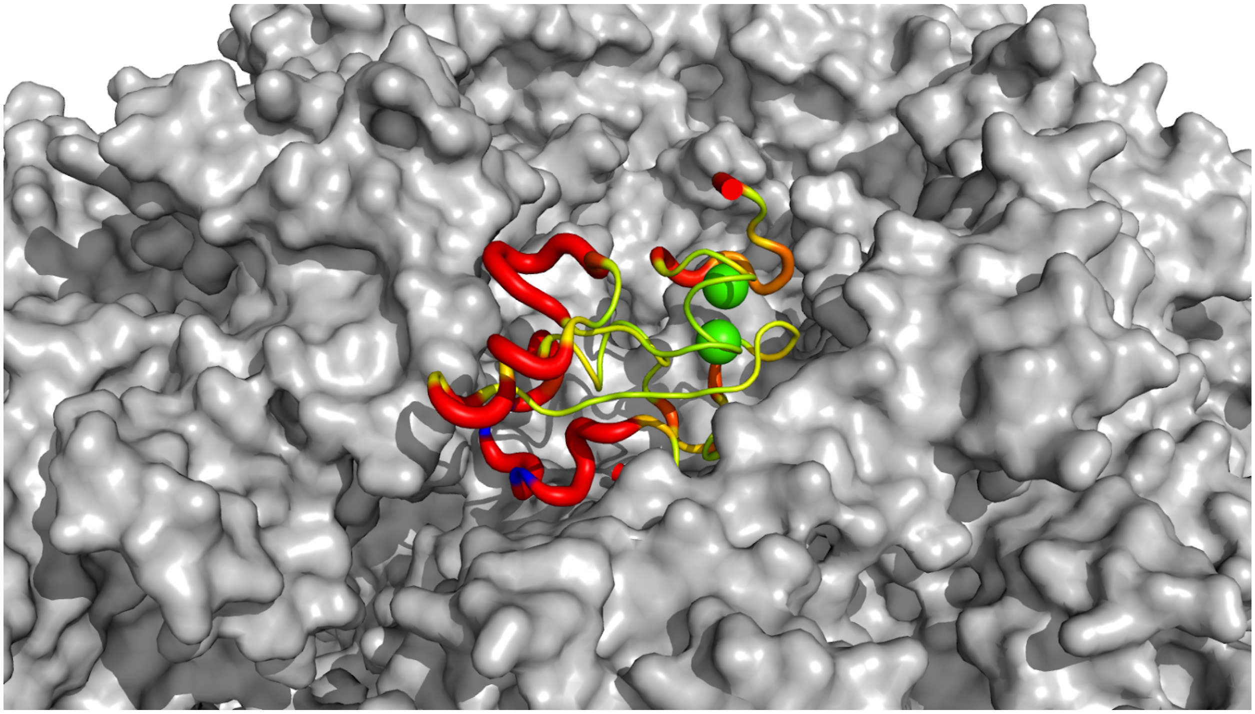

Figure S5. Single heptamerization 1 domain showing areas of structural differences with corresponding domain from the protective antigen (PA) of anthrax toxin. Thicker ribbon shows increased structural differences.

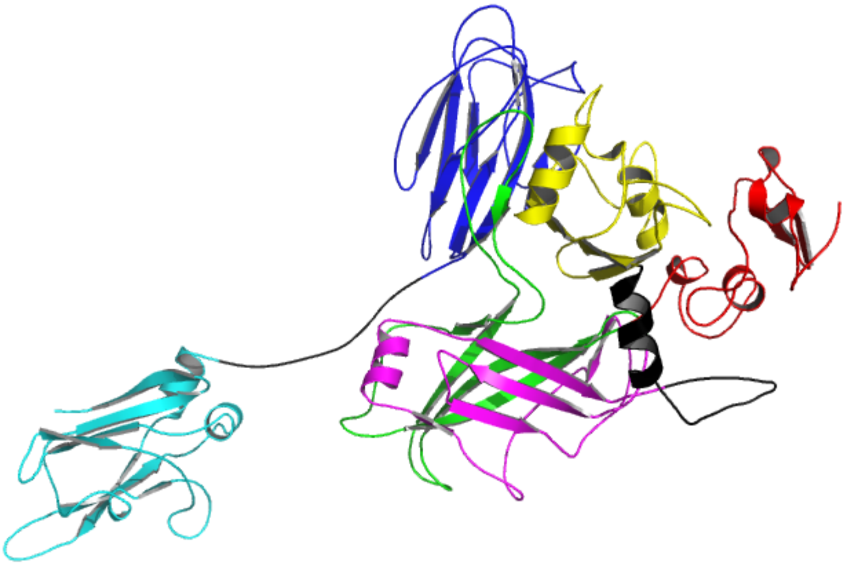

Figure S6. Overall structure of the CDTb subunit as it appears in the heptamer units of $^{Sym}$CDTb. The β-barrel domain (βBD) is shown in green, the second heptamerization domain (HD2) in magenta. The β-barrel domain packs against HD2 and its extended loop is placed in the crevice between the third heptamerization domain (HD3; yellow) and the first receptor binding domain (RBD1; blue).

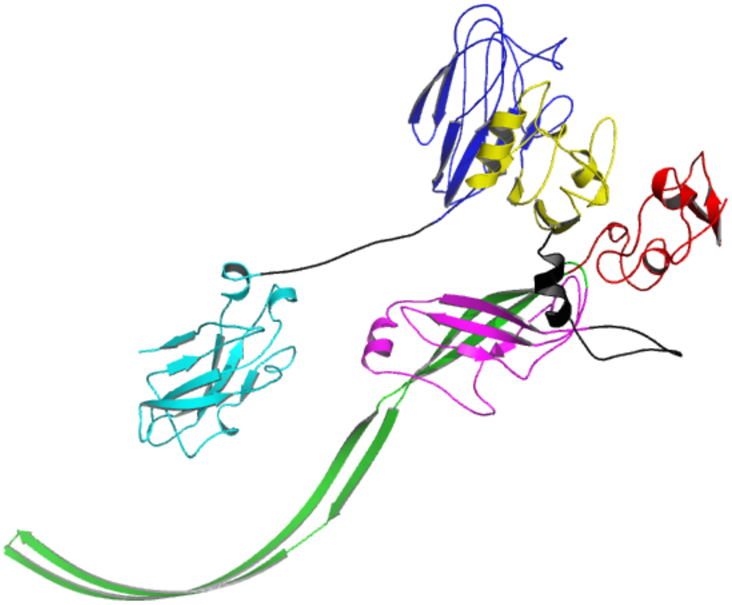

Figure S7. Overall structure of the CDTb subunit as it appears in the β-barrel heptamer units of $^{Asym}$CDTb. The βBD is shown in green.

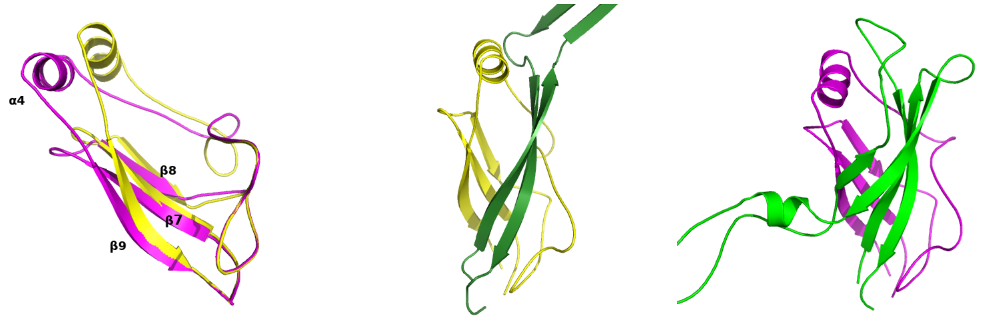


Figure S8. Comparison of the orientation(s) of HD2 in $^{Sym}$CDTb and $^{Asym}$CDTb. Left panel: superposition of HD2 in the “β-barrel heptamer” of $^{Asym}$CDTb (yellow) and the “non- β-barrel heptamer” of $^{Sym}$CDTb (magenta). Parts of the domain that contact the first and last β-strands of the βBD were aligned in this panel since those remain approximately in place between two forms of the domain. Middle panel: Packing of HD2 and βBD in the β-barrel heptamer of $^{Asym}$CDTb. Right panel: association of HD2 with βBD in the non-β-barrel heptamer of $^{Sym}$CDTb. Re-packing of the βBD causes shift in position of the α4 helix.

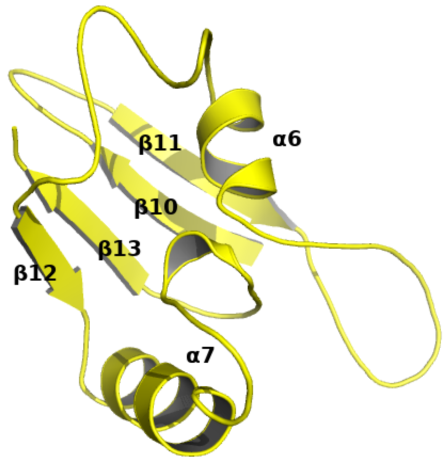


Figure S9.  Overall structure of the HD3.

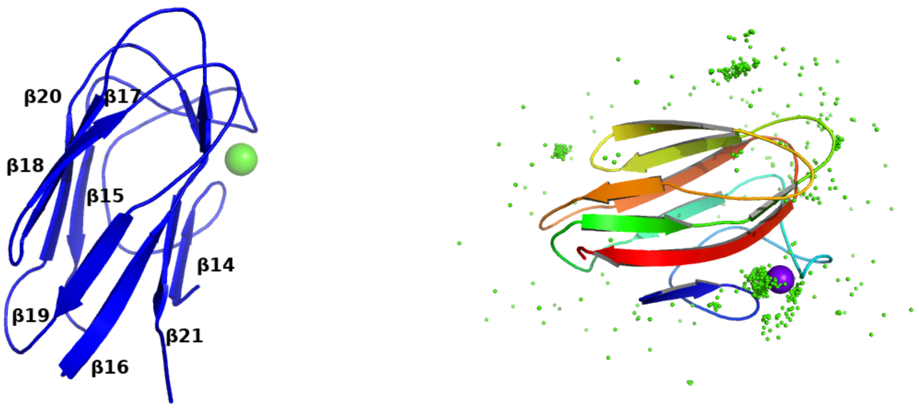


Figure S10. Overall structure of the RBD1 (left panel, calcium ion position shown in green). Right panel shows the location of calcium ions found in the structures of bacterial CBMs (green dots). Purple sphere shows the position of the calcium binding site found in RBD1. The secondary structure of RBD1 and the $Ca^{2+}$-dependence of folding, as determined by NMR (see **Fig. 6**) are in full agreement with the overall structure of RBD1 determined here via cryoEM.

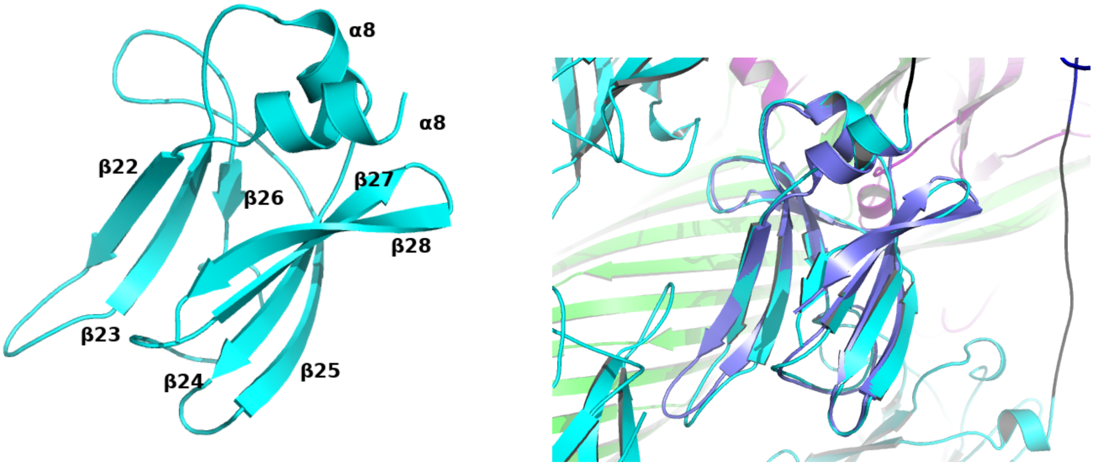


Figure S11. Overall structure of the RBD2 domain (left panel). Right panel shows superposition of the RBD2 crystal structure determined using an isolated domain (in purple) over the conformation found in the cryoEM model.

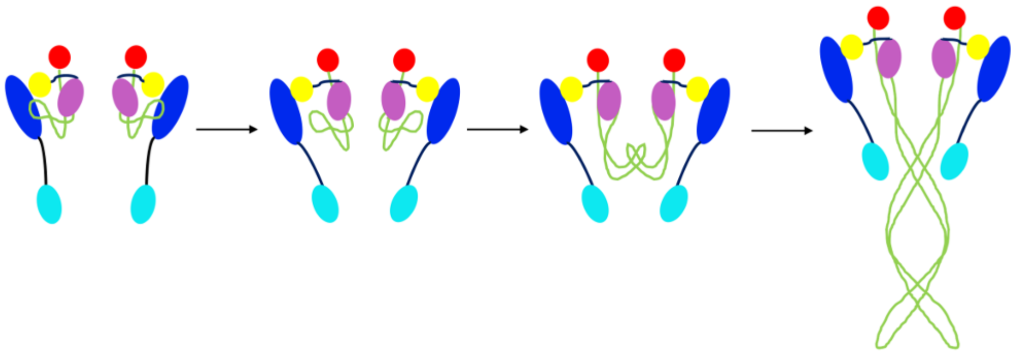

Figure S12. Illustration of possible conformational changes between the two conformations of CDTb heptamers as proposed by normal mode analysis calculations that may be related to its function. The color code is consistent with other figures with the heptamerization domain I (residues 212-297) shown in red, the beta-barrel domain (residues 298-401) in green wire, the heptamerization domain II (residues 402-486) in pink, the linker domain I (residues 487-513) in grey, the heptamerization domain III (residues 514-615) in yellow, the receptor binding domain I (residues 616-744) in blue, the linker domain II (residues 745-756) shown as purple wire and the receptor binding domain II (residues 757-876) in cyan.

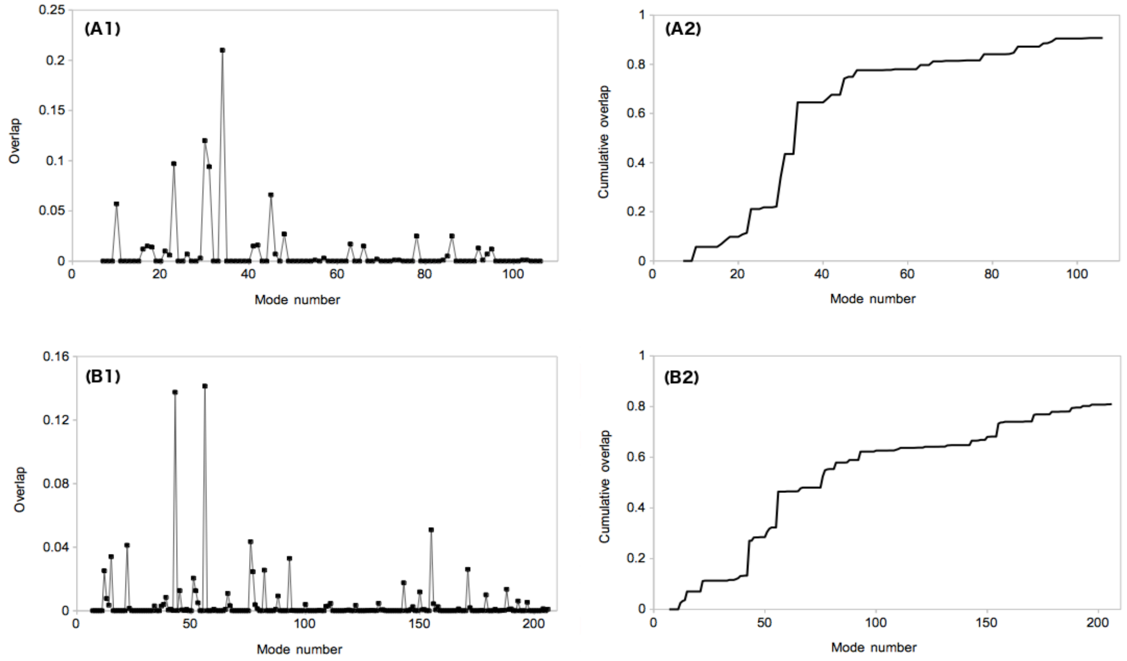


Figure S13. Overlap and cumulative overlap plots for normal modes calculated for heptamer (A) and 14mer (B) of CDTb that are related to the conformational change between the two conformations (from Figure S12).

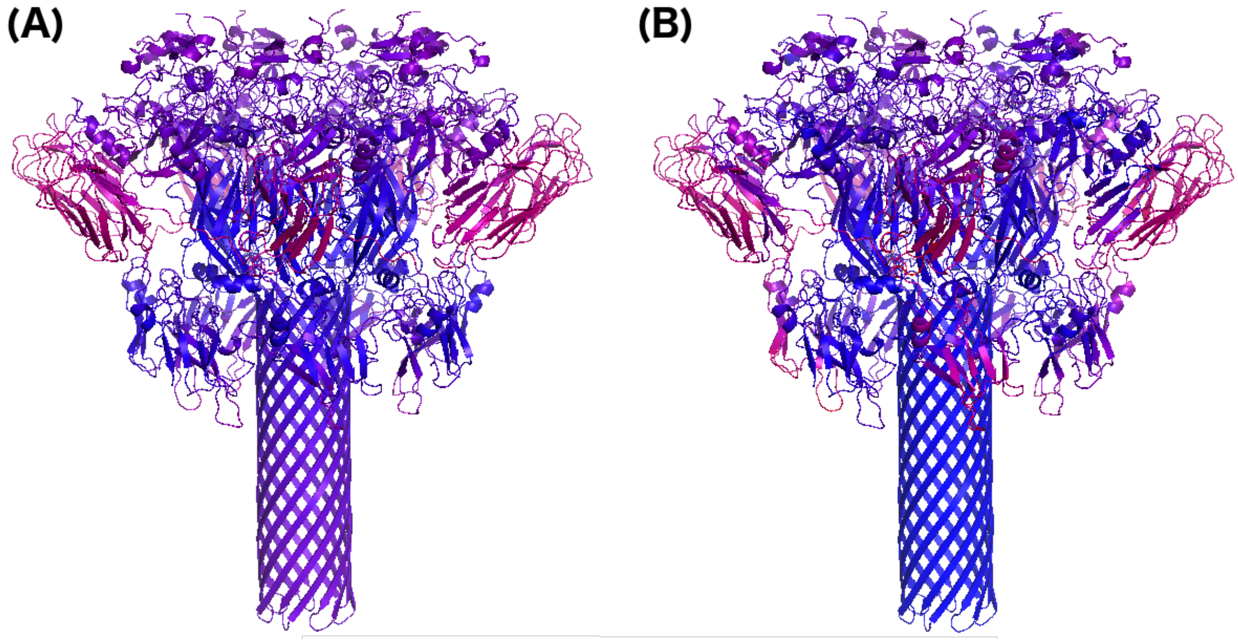


Figure S14. Two normal modes 34 and 30 (see Fig. S12) have the most overlap with the conformational change vectors and are predicted to be highly related to the conformation change between two conformations. RMSD mapped onto heptamer structures for mode 34 (A) and 30 (B) from Figure S12 with color scale in blue-purple-pink which means blue has the least RMSD while pink has the most RMSD. Of interest, it is observed that some of the largest RMSD values reside within RBD1, which is known to have a $Ca^{2+}$-dependent folding mechanism when isolated, as determined by NMR (See Fig. 6).
